# A new regulator of sporulation sheds light on spore morphogenesis and ballistospory in mushroom-forming fungi

**DOI:** 10.1101/2024.07.26.604922

**Authors:** Zhihao Hou, Zsolt Merényi, Yashu Yang, Yan Zhang, Árpád Csernetics, Balázs Bálint, Botond Hegedüs, Csenge Földi, Hongli Wu, Zsolt Kristóffy, Edit Ábrahám, Nikolett Miklovics, Máté Virágh, Xiao-Bin Liu, Nikolett Zsibrita, Zoltán Lipinszki, Ildiko Karcagi, Wei Gao, László G. Nagy

## Abstract

Sporulation is the most widespread means of reproduction and dispersal in fungi. In the Basidiomycota, sexual spores are produced on specialised cells known as basidia, from which they are discharged forcibly by a powered process called ballistospory, the highest known acceleration in nature. However, the genetics of sporulation, in particular postmeiotic events related to spore morphogenesis and ballistospory, remain poorly known. Here, we explore the genetics of these processes, based on a new, highly conserved transcription factor, Sporulation-Related Regulator 1 (SRR1), and its putative downstream regulatory network. Reverse genetics of *Srr1* in the model mushroom *Coprinopsis cinerea* and commercially produced oyster mushroom indicated a conserved role of *Srr1* in sporulation across Agaricomycetes. RNA-Seq analysis and motif-based inference of a hypothetical SRR1 gene regulatory network allowed delimiting putative targets regulated by SRR1 in a direct and indirect manner. Using this network and comparative genomics, we identified genes associated with ballistospory, including a putative SRR1-target chitinase, which was found to be required for normal spore production and morphology. Overall, our study offers new insights into the genetic mechanisms governing postmeiotic spore morphogenesis and ballistospory in the Agaricomycetes.

## Introduction

Sporulation is one of the most fundamental phases in the fungal life cycle. It is the primary means by which fungi reproduce, disperse to new habitats, survive unfavourable conditions and, in sexual spores, generate genetic variability for adaptation to new niches. Fungi produce massive amounts of spores, which can reach up to one billion/day for a single fruiting body^1^, that are carried by winds to new habitats. In mushroom-forming fungi (Agaricomycetes) air currents generated by the fruiting body can also contribute to lifting spores above ground so that they can travel significant distances from the parent organism^2^. *En masse*, spores pose health hazards and may even contribute to raindrop nucleation^3, 4^.

Spore formation includes the generation of haploid nuclei, the generation of spores and spore discharge. For the latter, fungi evolved unique mechanisms that ensure spores are shot forcibly at considerable distances from the reproductive cell, using lineage-specific mechanisms. In the Ascomycota this is achieved by building pressure in specialised cells called asci, whereas in the Basidiomycota, in which the spore discharge mechanism is known as ballistospory^5, 6^, spores are born externally on basidia. Ballistospory is a powered release mechanism during which the spore is catapulted from the basidia at initial accelerations exceeding 10,000 g, outpacing all other known rapid movements in nature^5^. The mechanism generating these ultrafast movements has only recently been elucidated^6^. While this improved our understanding of the physical laws of spore discharge, the genetics of spore formation and, in particular, of ballistospory remain unknown.

Beyond understanding fungal reproduction, understanding the genetics of spore production would benefit industrial applications and biomedicine too. Spores of commercially produced Agaricomycetes, such as *Pleurotus* spp. (oyster mushrooms) can harm technical instrumentation of production facilities^7^, can cause respiratory diseases known as mushroom worker’s lung, and can help strains escape cultivation to establish invasive populations or influence the genetic diversity of natural ones. As a result, there is a high demand for sporeless mushroom strains. Efforts to produce these strains in industrially relevant fungi^8^, primarily based on mutations in meiotic genes, have yielded industrially accepted sporeless alternatives of wild-type strains^9, 10, 11, 12^. However, for identifying novel target genes that can be used in strain improvements programs, uncovering the regulation of spore production is key.

In the Agaricomycetes spore morphogenesis is best understood in *Coprinopsis cinerea*^13^, though it has been studied in several other species too^14, 15, 16^. It starts after the completion of meiosis, which generates four haploid nuclei that will migrate into the developing spores. Spore morphogenesis has been divided into four stages and starts with the emergence of sterigmata, horn-like projections that serve as the platform for spore development^13^. In the next steps the spore first grows asymmetrically, then isodiametrically, paralleled by continuous changes to its wall structure and finally the abscission zone develops between the spore and the basidium. Morphogenesis concludes with forcible discharge, during which the spore is catapulted away from the basidium by forces generated by the coalescence of a water droplet (Buller’s drop) with the spore^5, 6^. Sporulation and ballistospory may have been one of the main constraints influencing the evolution of morphologies in the Basidiomycota^17^. The genetics of processes upstream of morphogenesis, such as meiotic events have been uncovered at great detail, particularly in *C. cinerea*, which has become a model species of meiosis^18, 19, 20, 21, 22, 23, 24^. However, the genetics of postmeiotic events, particularly morphogenesis and their regulation are virtually unknown.

In this study we identified a conserved transcription factor that regulates spore formation, which we term sporulation-related regulator, SRR1, a C2H2-type zinc-finger transcription factor. By reverse genetic analyses in *C. cinerea* and *Pleurotus cornucopiae* we show that SRR1 has a widely conserved role in regulating postmeiotic events of spore formation. RNA-Seq and motif analyses unveiled an SRR1 regulon of ∼600 genes, and revealed biochemical functions involved in spore formation. Genomic analyses identified genes that, based on expression and presence/absence patterns, are associated with the emergence and loss of ballistospory. This work uncovered a novel regulator of sporogenesis in Basidiomycota, its target genes and provided the first insights into the genes involved in spore morphogenesis and ballistospory.

## Results and Discussion

### *Srr1* is a novel regulator of sporulation in the Agaricomycetes

For identifying potential regulators of sporulation, we scrutinised genes upregulated in sporulating gill samples of the young fruiting body stage of *C. cinerea*^13^, compared to other tissues or other developmental stages, based on data from Krizsán et al. ^25^. We identified seven genes with an expression peak in gills, of which only one, *C. cinerea* 451915 was exclusively expressed in the gills, with no detectable expression (<10 CPM) in all other tissues and developmental stages (Supplementary Fig. 1A). *C. cinerea* 451915 also has the highest fold-change compared to the average expression in all stages (Supplementary Fig. 1B) and follows a gill-specific expression pattern in other basidiomycetes too, such as *Laccaria bicolor, Cyclocybe aegerita*, and *Pleurotus ostreatus* (Supplementary Fig. 1C-E) based on published data^26, 27, 28^.

We generated a knockout mutant for this gene in the self-compatible *C. cinerea* A43mutB43mut #326 pab1-1 strain, using a ribonucleoprotein (RNP) complex^29^. We targeted the locus with two RNPs, to generate a 1,666 base pair (bp) deletion (Supplementary Fig. 2B). The genotypes of the knockout mutant was confirmed by PCR using diagnostic primer pairs and RT-PCR to verify the lack of expression of the target gene (Supplementary Fig. 2C-D).

The 451915 knockout mutants displayed no phenotypic difference with the wild-type (WT) in the vegetative mycelium, oidiation capacity or early stages of fruiting body development (Fig. 1A and Supplementary Fig. 3A-B). Mature fruiting bodies, in contrast, had white caps, indicative of a failure to produce black spores (Fig. 1A and Supplementary Movie 1). Gills, cystidia and pseudopharaphyses did not differ from the WT and the autolytic process typical of *Coprinopsis* was also similar to that of WT (Fig. 1B-C). In the mutant, spore development is arrested at the stage when spore initials emerge at the sterigma tips (Fig. 1D), corresponding to stage one or two as defined in *C. cinerea*^13^. In the mutant these spore initials could not inflate further to reach a final spore shape and size. Fluorescent microscopy indicated that the basidia of the mutant contained four nuclei (Fig. 1D), indicative of successful completion of meiosis. Hereafter, we refer to *C. cinerea* 451915 as sporulation-related regulator 1, *Srr1*. *Srr1* encodes a 684 amino acid polypeptide which contains three C2H2-type zinc-finger domains (Supplementary Fig. 3C), indicating it is a transcription factor of the C2H2-type zinc finger family. Together these data indicate that *Srr1* mutants show a postmeiotic spore arrest in development, suggesting that SRR1 regulates morphogenetic processes following meiosis.

**Figure 1.**
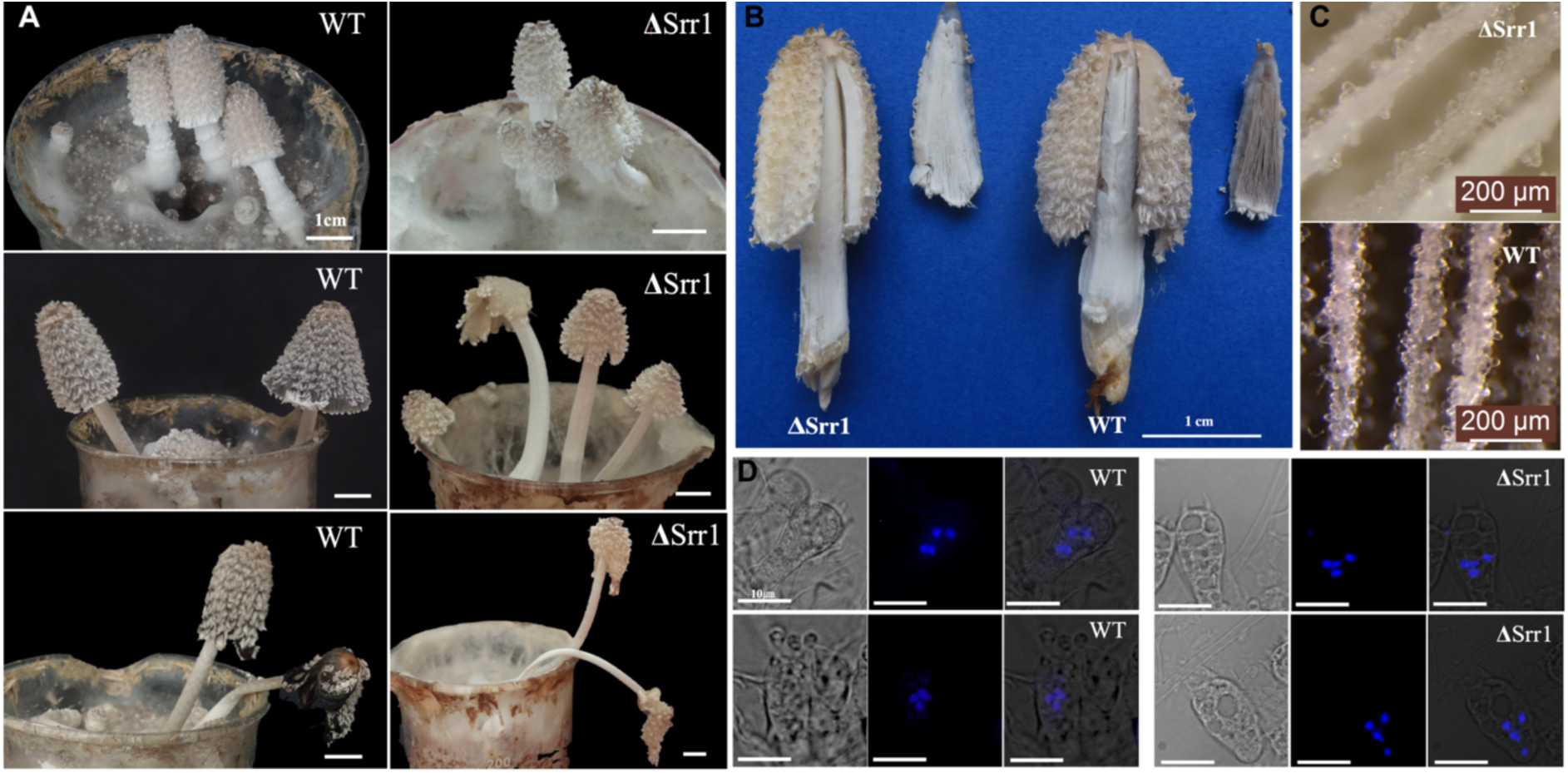
Phenotypic comparison of fruiting bodies, gills and basidia of ΔSrr1 and wild-type strains. A. Fruiting bodies at different developmental stages. B. Detailed phenotype of the fruiting bodies and the gills. C. Stereomicroscope images of cystidia and pseudopharaphyses. D. Fluorescent microscope images showing the number of nuclei in the basidia using Hoechst staining.

White cap mutations are indicative of failure in spore production in this species, due to the lack of spores that colour the caps black and include several classes that halt development at different stages^30^. *Srr1* mutations belong to the category that involves prespore formation, in which at least 13 genes have been reported using ultraviolet mutagenesis^31^, however, the corresponding genes have not been identified because of the randomness of the mutations. *Srr1* might be one of them and represents a novel transcription factor encoding gene that regulates spore formation.

### SRR1 is conserved across Agaricomycetes

To assess the distribution of SRR1 orthologs in the Agaricomycetes, we inferred a maximum likelihood gene tree for homologs of *C. cinerea* SRR1 identified in 189 whole genomes using blastp (E-value<10^−10^). Orthologs of SRR1 were defined as the most inclusive clade of the gene tree that does not comprises deep duplications. Our results suggest that SRR1 is widely conserved in the Agaricomycetes (Fig. 2A), with clear orthologs in the Agaricales, Boletales, Amylocorticiales, Polyporales, Jaapiales, Russulales and the Gloeophyllales, but missing in other Agaricomycetes orders (Fig. 2B). This suggests that SRR1 evolved after the Russulales split off from other Agaricomycetes. Within the Agaricales, the gene is conserved, except in the Schizophyllineae and Marasmiineae suborders, which lack orthologs due to gene losses. We found that SRR1 is a single-copy gene in most species that possess it, with few exceptions of recent duplications (e.g. *Suillus luteus*, see Fig. 2A).

**Figure 2.**
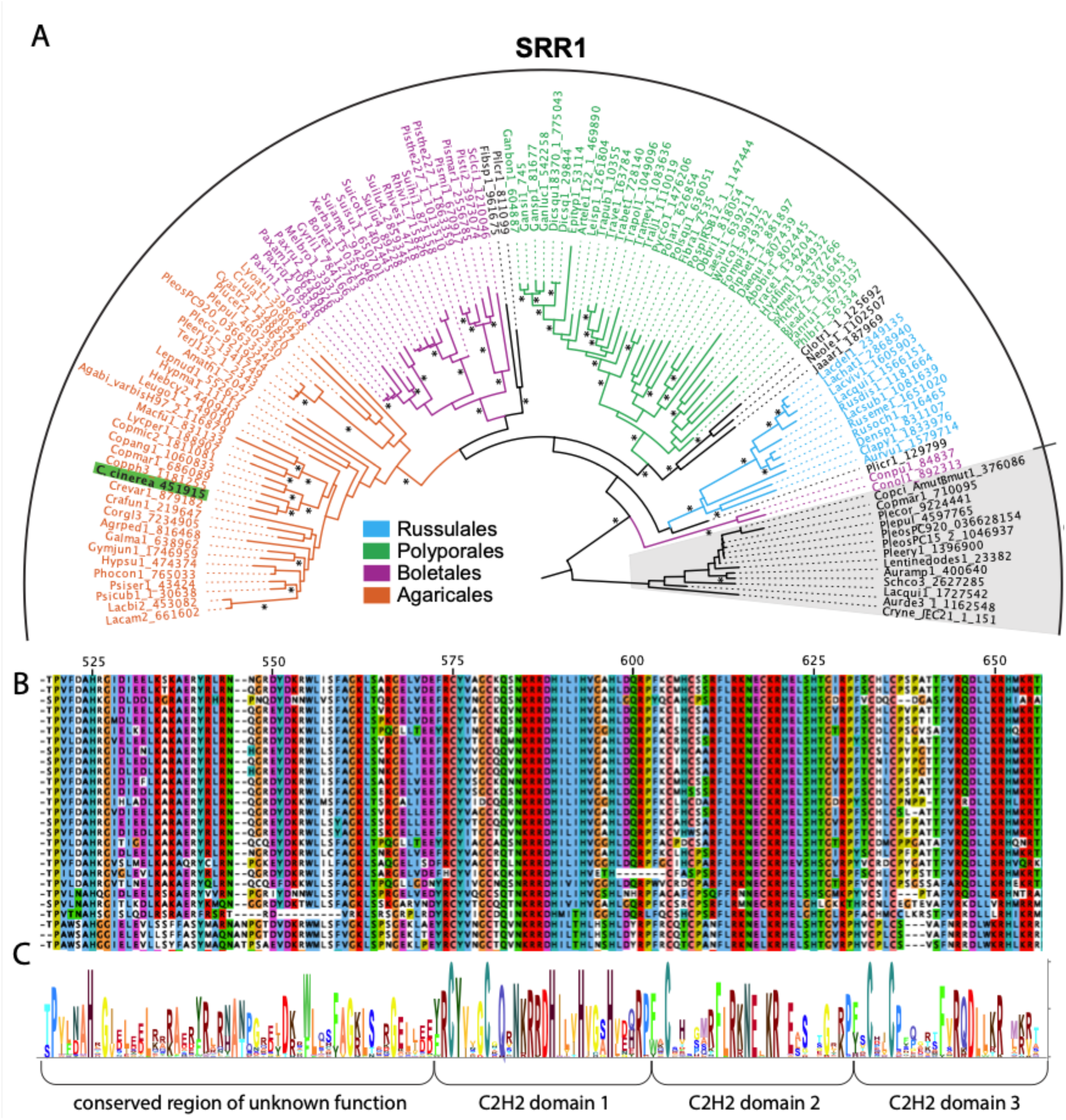
Evolutionary conservation of SRR1 across mushroom-forming fungi. A. maximum likelihood gene tree of SRR1 orthologs and homologs. Asterisk marks branches with >70% bootstrap support. B. the conservation of DNA binding domains and flanking regions of unknown function in the SRR1 polypeptide. Multiple sequence alignment shown only for the Agaricales. C. HMM logos of position-wise residue conservation in the DNA-binding domains of SRR1 orthologs.

The protein sequence shows overall low conservation, except for the three consecutive C2H2 type DNA-binding domain regions and a ∼50 amino acids (aa) upstream region, which are highly conserved and show a characteristic sequence signature that differs from that of other C2H2 families (Fig. 2C)

Outside Agaricomycetes, SRR1 did not have orthologs based on searches with the complete protein sequence. However, BLAST analysis of the DNA binding domain region (572 to 657 in the polypeptide) indicated a close similarity to Mld-1 of *Neurospora crassa*, which is involved in melanin production^32^.

### Conserved and overlapping role in oyster mushroom

The phylogenetic analyses of the SRR1 family revealed the presence of orthologs in the genus *Pleurotus*, which includes some of the most widely cultivated mushrooms worldwide. Based on previously published transcriptome data, the SRR1 orthologous gene of *P. ostreatus* was upregulated in gills^26^.

To analyse the role of the SRR1 orthologs in this genus, we used *P. cornucopiae* in which RNA interference (RNAi) has been established, to silence the g1654 gene (Plecor_9219244 in Fig. 2A). An RNAi-1654 vector was constructed, using the wild-type strain CCMSSC00406 as the host, and RNA interference transformation was carried out using *Agrobacterium*-mediated method. Quantitative PCR (qPCR) indicated that, compared to the wild-type strain, the expression level of g1654 in the gills of RNAi-g1654-4 and RNAi-g1654-14 strains decreased by 33.92% and 35.61%, respectively (Fig. 3A). These two strains were selected for subsequent experiments.

**Figure 3.**
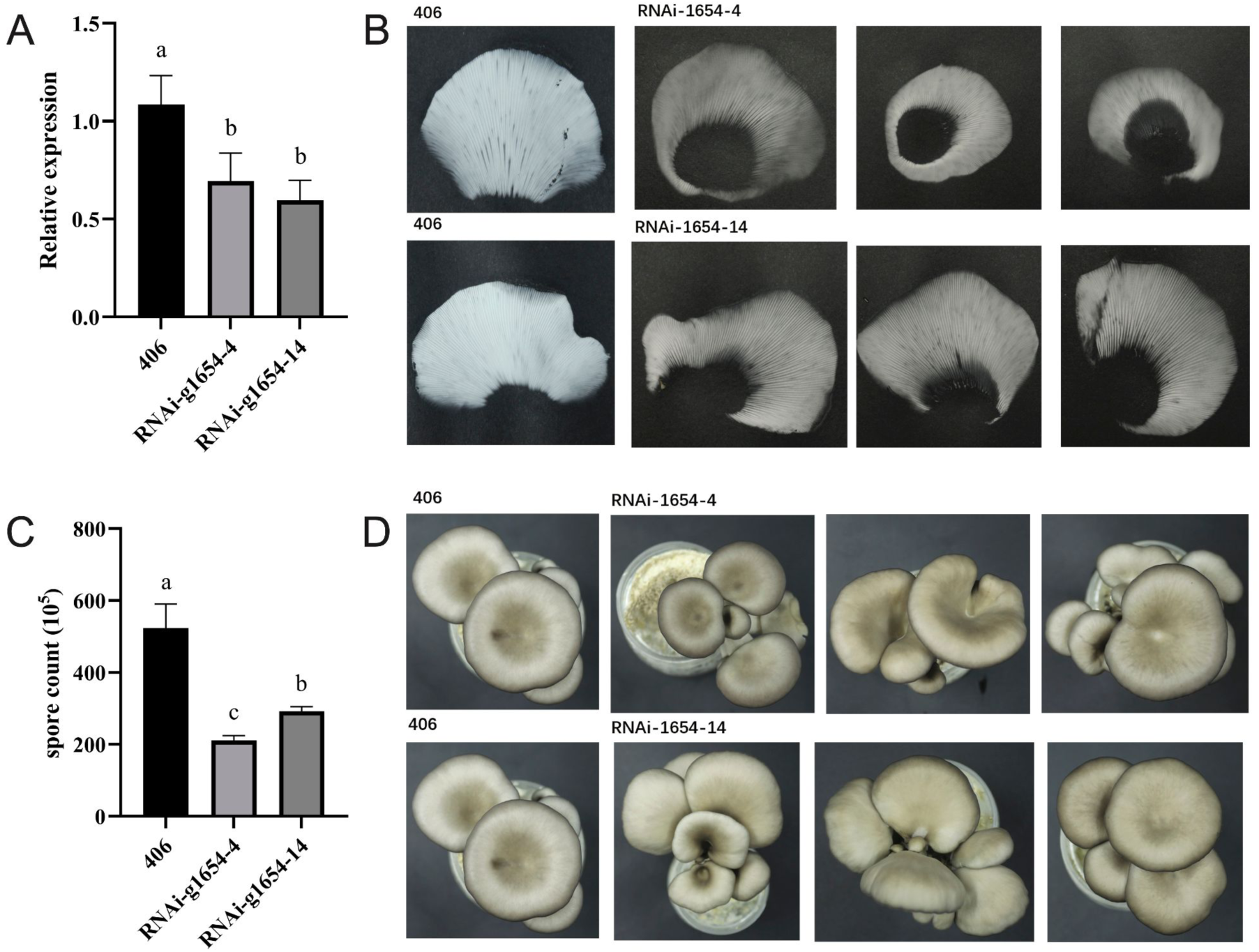
Comparison of P. cornucopiae Srr1 (g1654) expression, spore quantification and phenotype between silencing and wild-type (406) strains in P. cornucopiae. A. qPCR expression level in gills in WT and RNAi lines; B. spore prints of WT and the 1654-4 and 1654-4 RNAi lines; C. spore counts obtained from 0.5×0.5 cm cuttings of spore prints from (B) from WT and two RNAi lines. Significance groups are indicated by letters; D. Top view of fruiting body morphologies in WT and SRR1 RNAi strains.

Fruiting experiments were conducted with RNAi-g1654-4, RNAi-g1654-14, and the wild-type 406 strain. Compared to the wild-type strain, RNAi strains had nearly identical fruiting bodies (Fig. 3D). The spore prints of the RNAi strains were noticeably thinner and lighter in colour (Fig. 3B). In line with these results, spore quantity was significantly reduced in the RNAi-g1654-4 and RNAi-g1654-14 strains compared to the wild-type strain (Fig. 3C). These results suggest that g1654 plays a role in the spore production process of *P. cornucopiae*, and its reduced expression can explain the lower quantity of spores in RNAi strains.

### RNA-Seq reveals that SRR1 is mostly an activator

The sporulation process of *C. cinerea* Δ*Srr1* mutant stopped after the meiosis, when spore initials emerge at the sterigma tips. For understanding what genes SRR1 regulates, we sampled the gills of the WT and Δ*Srr1* at the time point when sporogenesis of Δ*Srr1* mutants stopped for RNA-Seq (Fig. 4A). Pearson correlation coefficients showed good grouping for both wild-type and Δ*Srr1* replicates (Fig. 4B). By analysing differentially expressed genes (DEGs, p≤0.05, |fold-change (FC)|≥2) we identified 154 and 559 genes that were significantly up- and downregulated, respectively in the Δ*Srr1* strain compared to WT (Fig. 4C and Supplementary Table 1a-b). The higher number of downregulated genes (559) compared to upregulated genes (154) suggests SRR1 is probably an activator, though DEGs likely include both direct and indirect targets. By cross-checking the expression level of downregulated DEGs with data from Krizsán et al.^25^, we found that most of them are highly expressed in the gill of young or mature fruiting bodies (Fig. 4D), indicating these genes become active in the gill during sporogenesis.

**Figure 4.**
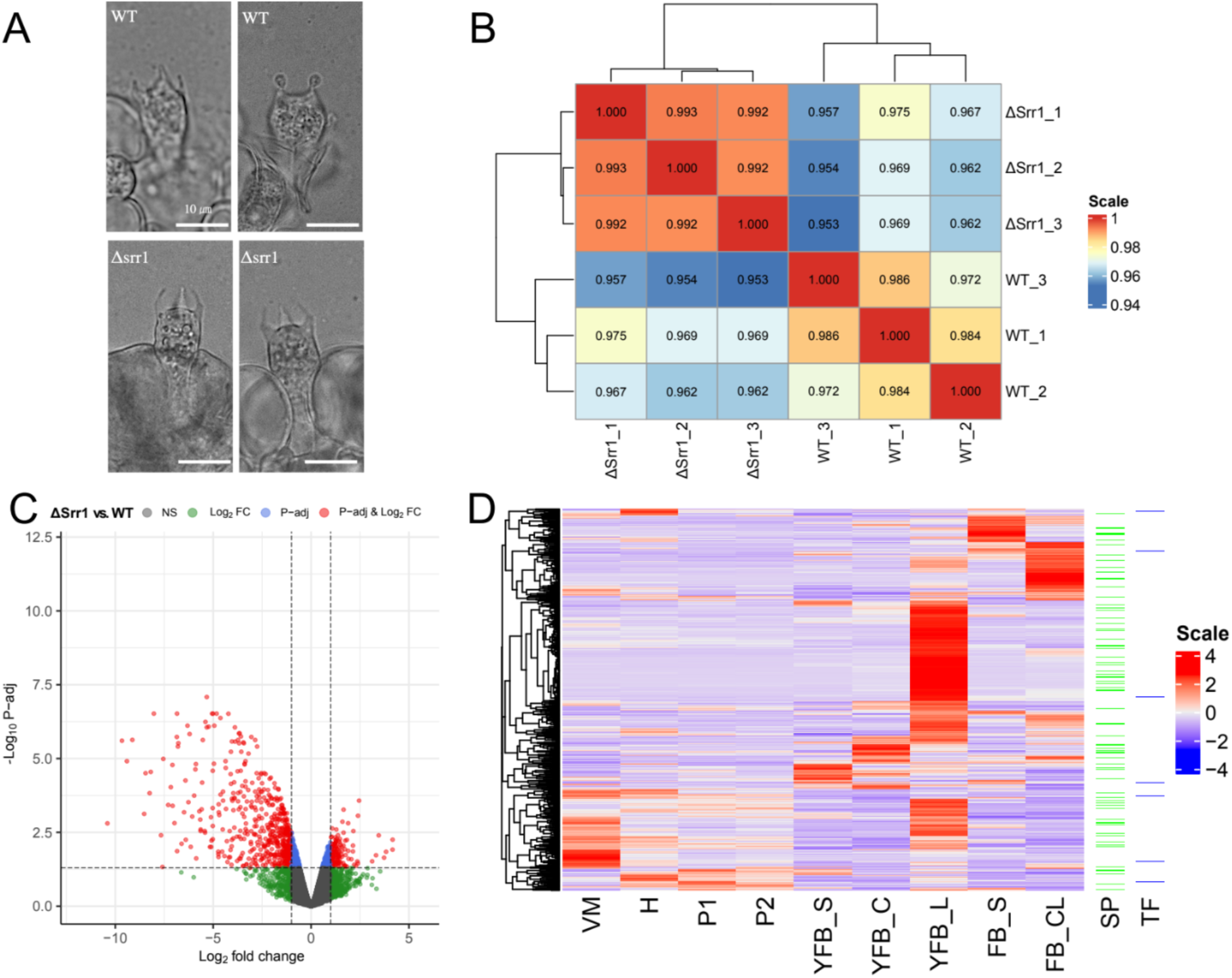
RNA-Seq analysis of ΔSrr1. A. sampling points of the ΔSrr1and mutant. B Heatmap of sample correlations. The numbers in the cell are the Pearson correlations across samples. C. Volcano plot of the differential expression genes. D. Heatmap of the expression of 599 FC_2 down-DEGs in Krizsán et al.’s dataset^25^, VM: vegetative mycelium, H: hyphal node, P1: primordia stage 1, P2: primordia stage 2, YFB_S: young fruiting body stipe, YFB_C: young fruiting body cap,YFB_L: young fruiting body lamella, FB: mature fruiting body stipe, FB_CL: mature fruiting body cap and lamella, SP: secreted protein, TF: transcription factor. Genes were clustered by Complete Linkage method based on Euclidean distance using the scaled Z-score of CPM values.

To characterise the functions of downregulated DEGs, we used GO and KEGG enrichment analysis. Overall, a significantly lower ratio of downregulated DEGs have any GO (Fisher’s Exact Test (FET), p-value=3.69×10^−12^-4.16×10^−04^) than all genes in the genome (Supplementary Table 2a), suggesting that the function of sporulation-related DEGs are particularly poorly known relative to the genome-wide average in *C. cinerea*. Nevertheless, we identified 103 significantly enriched GO terms (Hypergeometric test, p<0.05) (Supplementary Fig. 5 and Supplementary Table 2b). We found that enriched GO terms can be grouped into broad categories, corresponding to the fungal cell wall, amino acid biosynthesis, TORC signalling, sugar metabolism and the extracellular space. Hereafter, we discuss GO results based on this grouping (Supplementary Table 2b).

Terms related to the fungal cell wall (FCW) include aminoglycan metabolic process (GO:0006030, Hypergeometric test, p-value=1.46×10^−04^), cell wall chitin biosynthetic process (GO:0006038, Hypergeometric test, p-value=4.95×10^−02^) and 16 other related terms (Supplementary Table 2b). The cell wall consists of glucans, chitin and mannans, and sometimes lipids, whereas in addition the spore wall of *C. cinerea* contains melanin and has multiple layers which change dynamically during spore development^13, 33^. We here discuss FCW-related genes identified by GO and by manual annotation of *C. cinerea* cell wall genes^34^. Downregulated DEGs encode both cell wall synthetic and modifying enzymes (Supplementary Fig. 6).

There are two synthetic genes that encode chitin synthases (374670, logFC=-1.26 and 497826, logFC=-1.32) and one that encodes a Kre9/Knh1 family protein likely involved in β-1,6-glucan assembly (465604, logFC=-1.69) (Supplementary Fig. 6), with the highest expression in the young fruiting body gills (Supplementary Table 1b). On the other hand, among FCW-remodelling genes, we found four that are related to chitin, six to glucan and three to mannans (Supplementary Fig. 6). For chitin remodelling genes, two chitinases 543586 (logFC=-1.06) and 520359 (logFC=-2.50) were significantly downregulated in the Δ*Srr1*. They were named *ChiE1* and *ChiE2*, respectively in *C. cinerea*^35^, but were not functionally studied during fruiting body development in *C. cinerea*. Two putative chitin deacetylases, which convert chitin to chitosan (CE4, 360173, logFC=-2.13 and 42776, logFC=-1.06) were also downregulated in the Δ*Srr1*. Among putative glucan remodelling genes, three GH5 glucanases (375095, logFC=-5.05, 414471, logFC=-3.43; 503401, logFC= −4.04), two GH16 (449960, logFC=-1.16; 1000985, logFC=-3.61) one GH152 (500301, logFC=-2.51) are downregulated (Supplementary Table 1b). Moreover, we found a putative alginate lyase (543210, logFC=-3.73), which was hypothesized to have a relationship to the fungal cell wall (Supplementary Table 1b)^25^. As for the genes that might be related to the modification of the mannan layerlogFC=-5.53) is an α-mannosidase, while 143353 (GH2, logFC=-1.21) and 359841(GH5, logFC=-1.2) are β-mannosidases (Supplementary Table 1b). Taken together, these results suggest the fungal cell wall remodelling may be regulated by SRR1 during sporulation. Because Δ*Srr1* strains displayed normal deliquescence and gill cell morphology, we speculate that the downregulated fungal cell wall related genes are in fact involved in spore morphogenesis or abscission.

The next group of 22 enriched GO terms is linked mainly to aromatic amino acid (AAA) biosynthesis (Supplementary Table 2b and Supplementary Fig. 7). The KEGG pathway enrichment also shows that biosynthesis of amino acids (Hypergeometric test, p-value=2.26×10^−02^), and phenylalanine, tyrosine and tryptophan biosynthesis (Hypergeometric test, p-value=4.47×10^−05^) are significantly enriched among downregulated DEGs (Supplementary Table 2c). These GO terms correspond to seven genes, five of which (491583, 543926, 265487, 463372, 413668) participate in AAA biosynthesis, one (49306, logFC=-2.22) in histidine biosynthesis, and one (168151, logFC=-1.51) in arginine biosynthesis (Supplementary Fig. 7 and Supplementary Table 2d). Six of the seven genes are specifically upregulated in the young fruiting body gill (exception: 49306)^25^. AAAs are produced via a series of enzymatic reactions known as the shikimate pathway. 491583 (logFC=-2.38) is a 3-Deoxy-D-arabinoheptulosonate 7-phosphate (DAHP) synthase, the first enzyme in the shikimate pathway, which can control carbon input of the AAA biosynthesis (Supplementary Fig. 7)^36^. 543926 (logFC=-3.05) is a pentafunctional AROM polypeptide that catalyzes the formation of 5-enoyl-pyrovyl-shikimate-3-phosphate (Supplementary Fig. 7)^37^. 265487 (logFC=-3.56) is a chorismate synthase which catalyzes the production of chorismate, the common precursor for the synthesis of all three AAAs (phenylalanine, tyrosine and tryptophan) (Supplementary Fig. 7). 463372 (logFC=-1.33) and 413668 (logFC=-1.59) are the subunits of the anthranilate synthase complex that participate in tryptophan biosynthesis (Supplementary Fig. 7). These data suggest that the synthesis of AAAs is downregulated in the Δ*Srr1* strain. Because the synthesis of other amino acids seems less affected in Δ*Srr1* and as AAAs are the precursors of melanin synthesis which^38^, in *C. cinerea* happens only in spores, we hypothesise that their downregulation may be related to the lack of melanized black spores in the mutant.

The next functional category includes genes related to Target of Rapamycin (TOR) signalling (GO:0031929, Hypergeometric test, p-value=1.09×10^−02^), regulation of autophagy (GO:0010506, Hypergeometric test, p-value=3.02×10^−02^) and five related terms (Supplementary Table 2b). We found three *C. cinerea* genes in this category, all of which act in or upstream of the TOR pathway, a conserved nutritional signalling cascade (Supplementary Fig. 8A-B). Two genes (498449, logFC=-1.06 and 470617, logFC=-1.43) are upstream of the TORC1 complex, while one (495775, logFC=-1.83) is part of TORC2. Of the two genes upstream of TORC1, 470617 might belong to the TORC1-activating SEACAT complex, based on sequence similarity, whereas 498449 belong to the TORC1-inhibiting SEACIT complex (Supplementary Fig. 8A)^39^. Regarding TORC2, 495775 is the ortholog of *S. cerevisiae* TSC11/AVO3 which is an essential subunit of TORC2 complex (Supplementary Fig. 8B). As TORC1 and TORC2 are involved in the regulation of diverse cellular processes (including autophagy and cytokinesis in yeasts) and responses to nutrient availability (e.g. amino acids), it is hard to speculate what their downregulation in our data means for sporogenesis. However, considering the strong signal for amino acid biosynthesis among downregulated DEGs, it is possible that the downregulation of TOR pathway components may be related to the amino acid deficiency that conceivably emerges in the Δ*Srr1* mutant. Alternatively, during yeast cytokinesis the TOR2 complex regulates actomyosin ring constriction, which may also be relevant during sporulation.

The enrichment of the term extracellular region (GO:0005576, Hypergeometric test, p-value=4.07×10^−09^) (Supplementary Table 2b) suggests that extracellular and/or secreted proteins may be particularly affected in the Δ*Srr1* strain. Therefore, we checked the secreted proteins among downregulated DEGs. We found that downregulated DEGs (15.74%, 88/559) that encode secreted proteins are significantly enriched (FET, p-value=4.74 x 10^−07^) among DEGs compared to the whole genome (8.96%, 1220/13615) (Fig. 4D and Supplementary Table 1b). DEGs encoding secreted proteins covered diverse functions, including predicted peptidases, fasciclins, cutinases, putatively glucan-related cellulose-binding domains (CBM1), and lysozymes, among others (Supplementary Table 1b). Previous studies revealed that the endoplasmic reticulum (ER) and Golgi vesicles were becoming active during spore formation in *C. cinerea*^13, 40^. This, combined with our results suggest an increased biosynthesis of secreted proteins during sporulation, potentially to fulfil functions in adhesion, cell wall remodelling or defence, among others.

Finally, we identified GO terms related to carbohydrate metabolic processes which correspond to malic enzymes of *C. cinerea*. All three malic enzymes (380777, logFC=-2.40; 273609, logFC-=-1.97; 497233, logFC=-1.14) encoded by the genome of this species were downregulated in the Δ*Srr1* strain. While this appears to be a strong signal, its relationship to sporulation is currently obscure, as malic enzymes catalyse the oxidative decarboxylation of malate to pyruvate and, as such, are involved in diverse primary metabolic processes.

To approximate the downstream transcriptional network of SRR1, we identified transcriptional regulators downregulated in the mutant and found seven genes encoding transcription factors (Fig. 4D). Four of them are C2H2-type Zinc finger transcription factors including 447714 (logFC=-3.93), 537559 (logFC=-5.11), 377260 (logFC=-1.32) and 467166 (ogFC=-1.90). 188021 (logFC=-1.29) belongs to the High Mobility Group Box family. 438653 (logFC=-1.57) is a member of the Gti1/Pac2 family, and 541612 (logFC=-1.02) contains the TEA/ATTS domain. Based on developmental expression data, 447714 has a peak expression in young fruiting body gills (Supplementary Table 1b), and 188021 is upregulated in the mature fruiting body gill indicating potential links to sporulation.

### Motif and network analyses highlight putative target genes of SRR1

To better understand the regulation of SRR1, we performed motif analyses of downregulated genes based on six alternative promoter datasets. Considering the uncertainty in defining *C. cinerea* promoters, and regulatory functions of promoter-proximal introns^41^, we first created six promoter datasets, with different lengths (300 bp, 500 bp and intergenic region with first intron) upstream of the start codon for both fold-change two (FC2, 558 genes) and fold-change four (FC4, 266 genes) downregulated genes (excluding *Srr1*) (Fig. 5A). For investigating the binding motif of SRR1, we inferred motifs using STREME for these six datasets. We obtained significantly enriched (p-value<0.05, Binomial test) motif(s) for each of the tested datasets with variants of one motif, GTGGCTGAC, appearing in all six (Fig 5B and Supplementary Table 3a). This motif was found in the promoters of 106-260 (27.06-46.99%) genes among different datasets (Fig. 5B and Supplementary Table 3b). We obtained a similar motif (GTGGCDHWG) by an independent predictor^42^, which only uses information from the protein sequence of SRR1(Fig. 5C).

**Figure 5.**
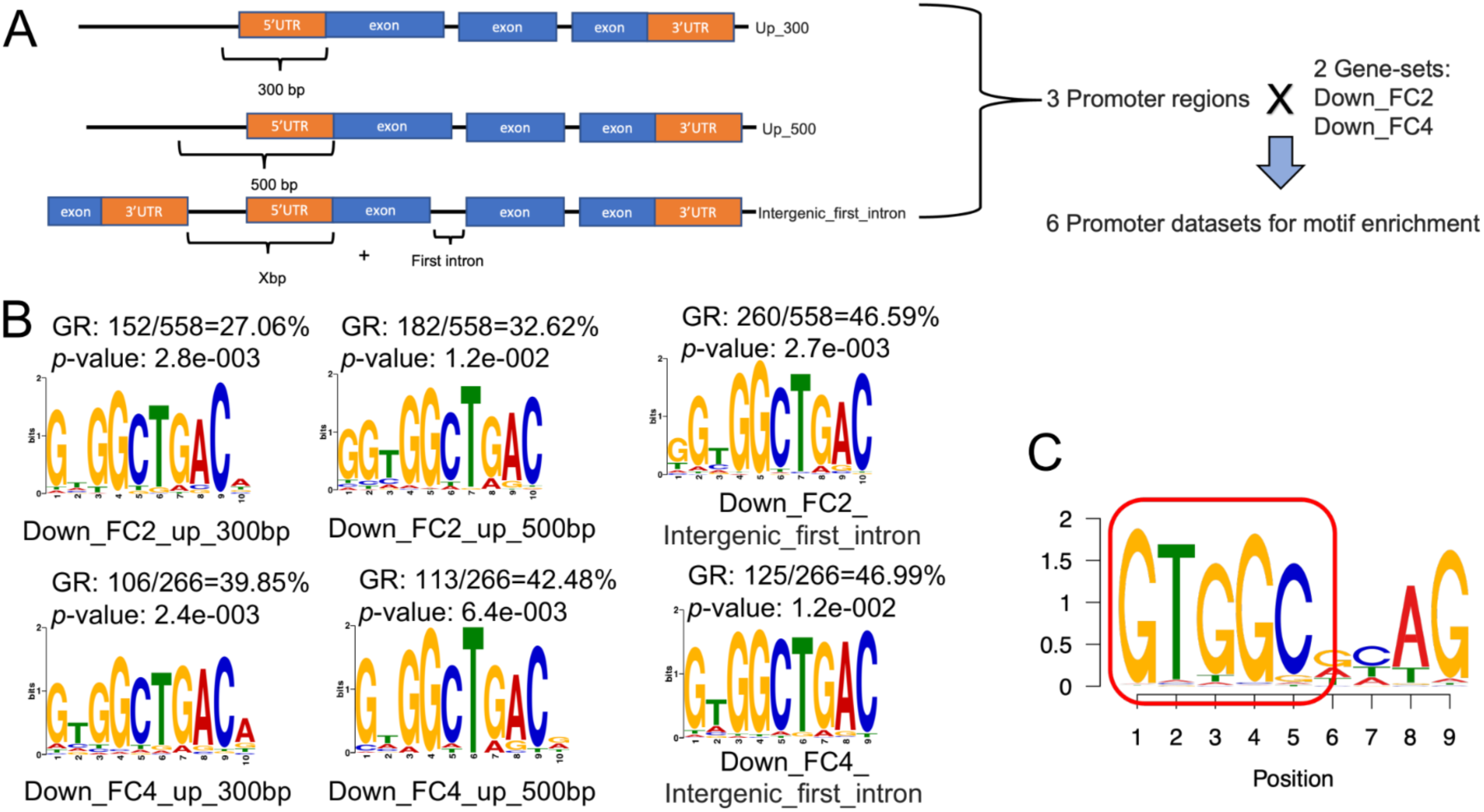
Motif analysis of the downregulated DEGs. A. The diagram of the generation of promoter datasets. B. The inferred motifs among different promoter datasets. GR: Gene ratio, refers to the genes with inferred motif/all the genes in the datasets, p-value refers to the Binomial test result for the enrichment. C. The prediction with DNA binding site predictor for Cys2His2 Zinc Finger Proteins. Red rectangle refers to the motif that has a high similarity with the inferred motifs in panel B.

For further analyses we chose the dataset comprising 558 downregulated genes (FC≥2), the whole intergenic region upstream of the gene plus the first intron (Down_FC2_Intergenic_first_intron). In this dataset, 260 of the 558 genes contained at least one occurrence of the GTGGCTGAC variants. The distribution of the inferred motif in the putative promoters is shown in Supplementary Fig. 9.

For understanding the gene regulatory network of SRR1, we compiled available evidence of potential upstream and downstream factors, target genes and inferred regulatory relationship into a hypothetical gene regulatory network (Fig. 6). In addition to *Srr1*, we identified six other gill-upregulated TFs (Supplementary Fig. 1A). Because none of them were differentially expressed in the Δ*Srr1* strain, we hypothesise some or all of them possibly act as upstream regulator(s) of *Srr1*, in combination with other signals (Fig. 6). It is possible that activation of *Srr1* is linked to the completion of cell cycle events after meiosis. Downstream of SRR1, our motif analyses identified two downregulated TFs (447714 and 467166) that have the predicted SRR1 binding motif in their promoters and are thus putative direct targets of SRR1(Supplementary Table 3b). Another five downregulated TF genes had no SRR1 motif in their promoters, hence we infer they are not directly regulated by SRR1. Both direct and non-direct TFs may regulate the downstream genes, including those associated with fungal cell wall, aromatic amino acid synthesis, secreted proteins and TOR signalling (Fig. 6).

**Figure 6.**
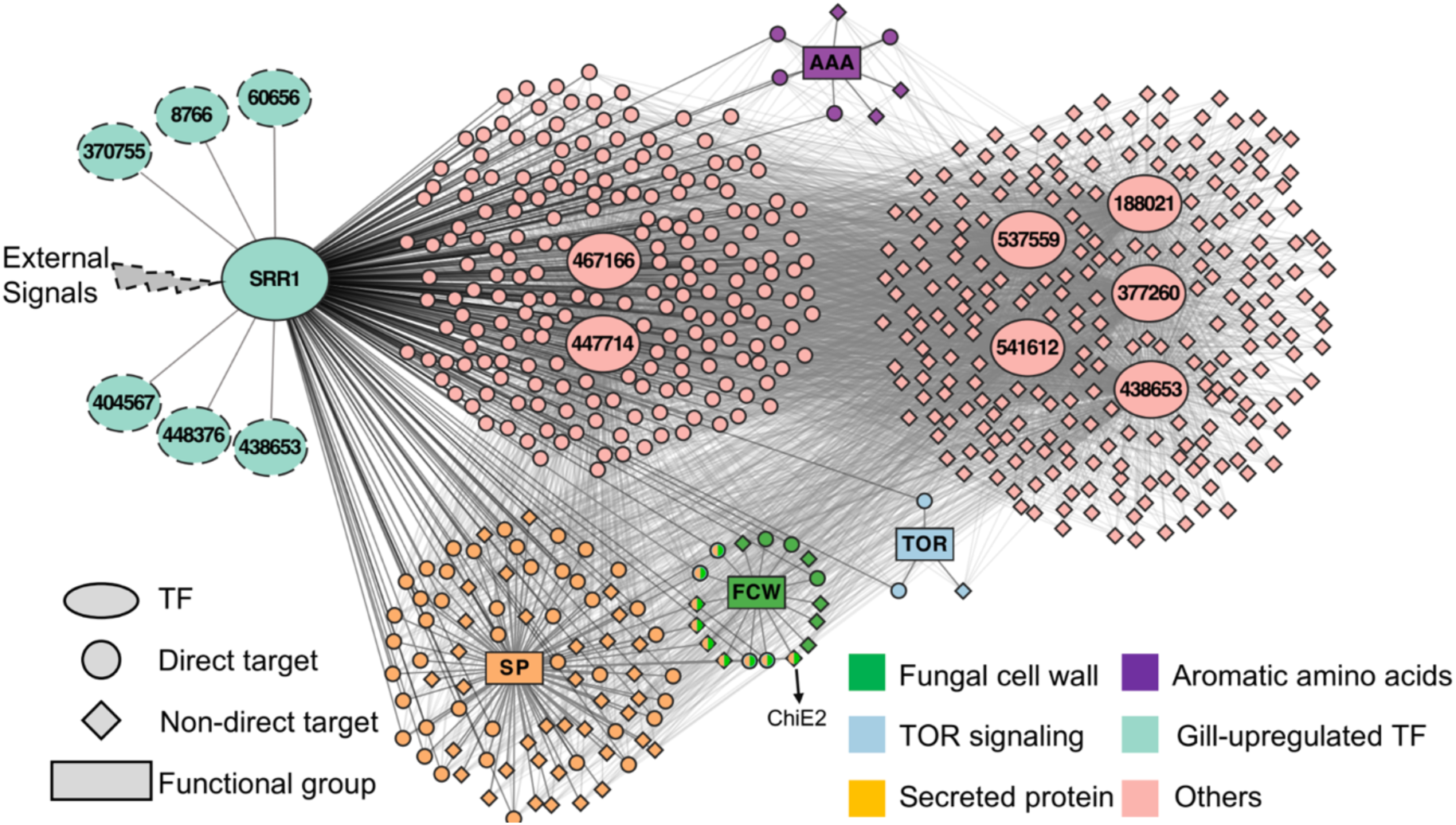
The hypothetical gene regulatory network of SRR1. The shapes represent the category annotation and the colours represent different functional groups. The dashed lines of the upstream transcription factors represent predictions based only on the gill-upregulated expression pattern.

Taken together, our analyses provided a clear signal for the putative binding motif of SRR1 and allowed approximating its regulon in *C. cinerea*. By integrating the available information, we propose a hypothetical regulatory network in which SRR1 acts as a key factor triggered by upstream TFs or signals and regulates the downstream targets to participate in sporogenesis and ballistospory.

### Comparative genomics of putative SRR1 targets reveals a link to ballistospory

Ballistospory is a highly streamlined basidiomycete process that involves the highest acceleration recorded in nature so far^6^. Here we attempted to uncover the genetic background of ballistospory by comparative genomics of putative SRR1 target genes. Since the SRR1 mutant is arrested at the start of spore morphogenesis, we hypothesize that genes responsible for ballistospory should also be among direct or downstream SRR1 targets. To address this, we analysed a 189-genome dataset using two approaches. First, using phylostratigraphy, we asked if evolutionary origins of downregulated genes in the Δ*Srr1* strain globally coincide with that of ballistospory. Second, we identified gene orthogroups whose presence/absence across basidiomycetes correlates with that of ballistospory, and we checked whether these candidates overlap with putative direct target genes of SRR1.

For the phylostratigraphic analysis of downregulated DEGs, we compiled a 189-genome dataset, comprising 164 ballistosporic species, and 18 that lost ballistospory (henceforth secondarily non-ballistosporic), such as puffballs, bird’s nest and false truffles, as well as seven Ascomycota as outgroup. The evolutionary origins of each of the 559 downregulated DEGs of *C. cinerea* were inferred by taking the most recent common ancestor of species possessing orthologs (Fig. 7). We found that the origin of 50 genes (9%) coincide with that of ballistospory in the Basidiomycota (including ChiE2), whereas most genes are either more ancient (152 genes, 27%) or specific to the genus *Coprinopsis* (120 genes, 21.5%), with 42.4% of genes originating in between. The age distribution of genes thus obtained strongly resembles the usual phylostratigraphic pattern in Agaricomycotina genomes^26^.Compared to all *C. cinerea* genes, we found that downregulated DEGs are significantly underrepresented in the most ancient phylostratum (FET, p-value=4.2×10^−11^), suggesting that the direct target genes of SRR1 are mainly Basidiomycota-specific. Nevertheless, we failed to detect any enrichment of SRR1 DEGs in the node where ballistospory might have emerged (FET, p-value=0.58), whereas three nodes showed significant enrichment of target genes (FET, p<0.05). This may indicate that genes required for ballistospory are not in high numbers or that they evolved earlier or later during evolution.

**Figure 7.**
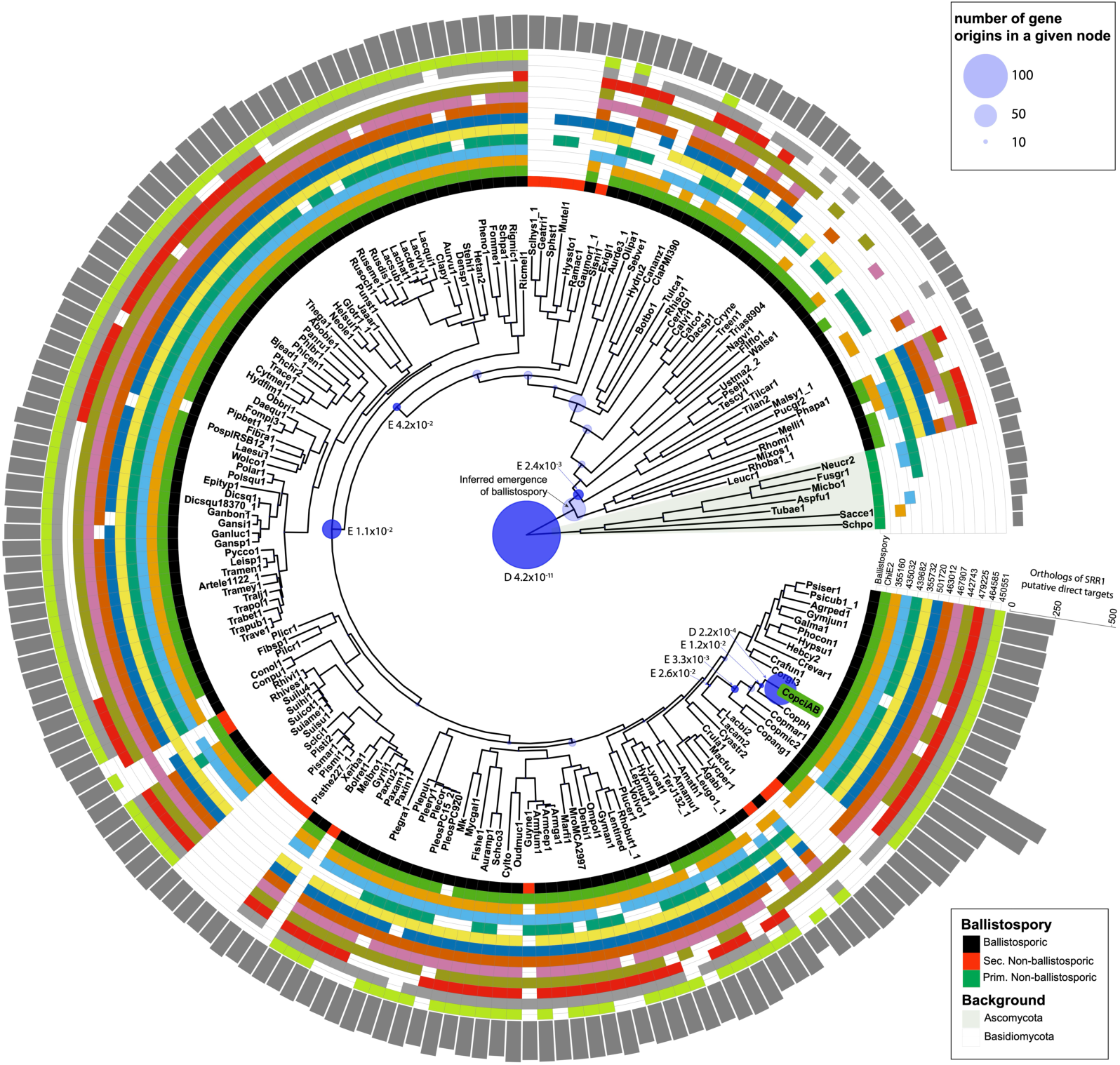
Phylostratigraphic analysis for putative direct target genes of SRR1 and the identification of candidate genes related to ballistospory. The species tree depicts the 189 species used in the analyses. The occurrence of ballistospory, and its independent losses in secondarily non-ballistosporic species (18 species in 8 independent lineages in our dataset) are shown by the innermost track with black and red, respectively. Phylostratigraphic analysis of downregulated DEGs in the ΔSrr1 strain (blue bubbles) illustrate the enrichment of gene origins in Basidiomycota nodes. Twelve candidate ballistospory-related genes are shown by coloured tracks (white means missing), these are mostly absent in non-ballistosporic lineages, but conserved among other Basidiomycota. Significant and non-significant enrichment or depletion are marked with opaque and transparent blue bubbles, respectively. ‘E’ or ‘D’ next to p-value (FET), stand for enrichment and depletion. The grey bar graph shows the number of orthologs of SRR1 DEGs present in each species. Explanation of species names can be found in Supplementary Table 4.

In our second approach, we aimed to identify ballistospory-related genes independently of the SRR1 targets. We hypothesised that the genes responsible for ballistospory may have been lost in some or all of the eight secondarily non-ballistosporic lineages. We searched for genes conserved in >50% of ballistosporic species, lost in at least five secondarily non-ballistosporic lineages and that show conserved lamellae-specific expression based on previous studies^25, 43^. Using this strategy, we identified twelve genes (Fig. 7 and Supplementary Table 5), most of which encode proteins with only generic or no predicted function, such as a nucleotide-sugar transferase, a peptidase, a serine-threonine exporter, and a GH25 lysozyme, among others. We also found two genes related to the fungal cell wall, a putative expansin (467907) and the GH18 family chitinase ChiE2 (520359). *ChiE2* was downregulated in the Δ*Srr1* strain, though it does not seem to be a direct target of SRR1. This gene and its orthologs showed the highest expression in the lamellae of *C. cinerea*, *Armillaria ostoyae*, *Lentinus tigrinus* and *P. ostreatus* and caps of (*L. bicolor* and *Lentinula edodes* in which dataset lamellae samples were not available), consistent with a narrow role in sporulation^43^. Phylogenetic analyses of ChiE2 orthologs and related chitinases confirmed the phylostratigraphic and gene loss patterns for this gene (Supplementary Fig. 10).

To test if *ChiE2* is involved in ballistospory, we generated knockout mutants in *C. cinerea.* The Δ*ChiE2* strains showed normal fruiting body development, including proper autolysis, and normal basidia with curved sterigmata, but produced spore-poor caps compared to the WT (Fig. 8A-C and Supplementary Movie 2). The Δ*ChiE2* strain produced asymmetrical spores that had a lighter colour and less consistent shape than those of WT (Fig. 8C), suggesting spore inflation, maturation or shape problems. In line with macroscopic observations, Δ*ChiE2* had a significantly lower number of basidiospores than the WT (Fig. 8D). The length and aspect ratio of the spores were also different and more variable in the Δ*ChiE2* strain than in the WT strain, further underscoring the sporulation-associated defect caused by its deletion (Fig. 8D). Because autolysis and powered spore discharge happen simultaneously in *C. cinerea*, observing the latter is challenging. Therefore, we cannot conclude with certainty if the *ChiE2* knockout affected ballistospory directly. Nevertheless, two prerequisites of ballistospory, spore asymmetry and curved sterigmata were present in the mutant^17, 44^, although spore shape, size and aspect ratio showed large variability, which might affect spore discharge. Spore shape is one of the important determinants of the maximum size of the Buller’s drop, which affects the launch energy during the spore discharge^45^. Considering all above, the results suggest *ChiE2* plays an important role in sporulation, maybe by participating in spore wall assembly/remodelling to result in spore shaping which facilitates the ballistospory. Since the ballistospory mechanism is still not fully clear, and autolysis happens almost simultaneously with ballistospory in *C. cinerea*, more investigation is needed for understanding of the role of *ChiE2* in ballistospory.

**Figure 8.**
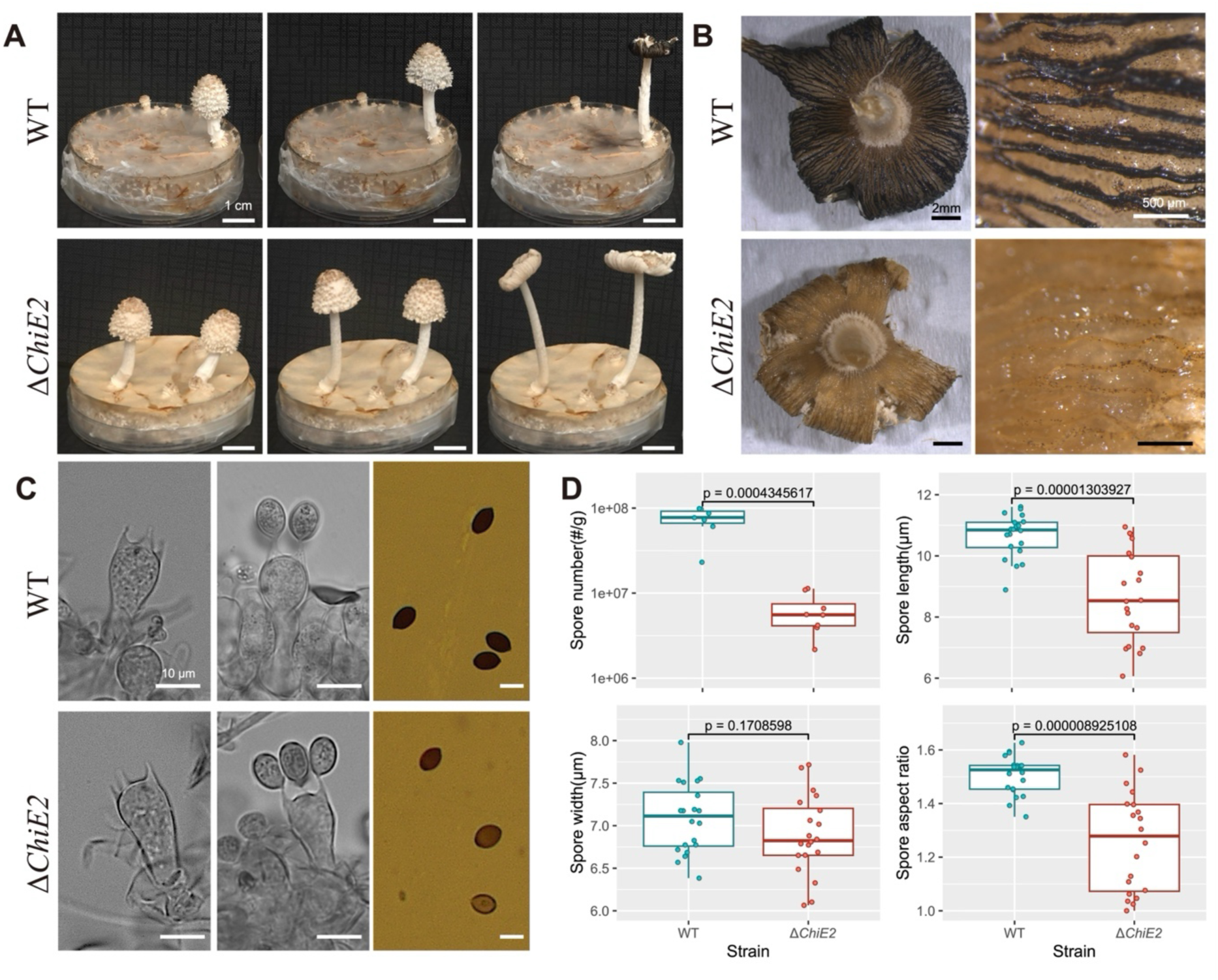
Comparison between ΔChiE2 and wild-type strains, including phenotype of fruiting bodies, gills, basidia and basidiospores, along with quantitative comparisons of the amount and shape of basidiospores. A. Fruiting body phenotype during developmental stages preceding sporulation. B. Stereomicroscope images of autolysing gills. C. Basidial morphology and mature spores. D. Quantitative comparisons for the basidiospores. Boxplot showing, clockwise, the number of spores per unit mass of cap, the distributions of basidiospore length, width, and aspect ratio (length divided by width). Significance was assessed by Student’s t test.

## Conclusions

In this study we described a novel transcriptional regulator (*Srr1*) and a gene network associated with postmeiotic events of sporulation, including spore morphogenesis and forcible spore discharge. We inferred its putative direct and indirect target genes in the model mushroom *C. cinerea* and approximated the regulatory network orchestrating spore morphogenesis. To identify genes linked to ballistospory, we combined comparative genomics, phylostratigraphy and the SRR1 regulatory network, and shortlisted several candidates, one of which was further analysed by reverse genetics.

We identified *Srr1* based on its nearly exclusive expression in gills of pre-sporulation fruiting bodies. SRR1 is a transcription factor with three C2H2-type zinc finger domains, and is conserved across the Agaricomycetes. The *Srr1* knockout mutant produced healthy fruiting bodies without spores, due to a postmeiotic arrest of sporogenesis after the emergence of spore initials at the sterigma tips (stage 2)^13^. Similar observations made in *Pleurotus cornucopiae* indicating that the role of *Srr1* is conserved. This phenotype corresponds to what has previously been reported as white cap or sporeless mutants in *C. cinerea*^19, 30, 31^. Most such mutations affect meiotic or DNA-repair genes (reviewed in Kües^40^), whereas SRR1 affects the postmeiotic process, which opened the way for dissecting morphogenetic aspects of sporulation.

Sporeless mutants have been generated in multiple agaricomycete species, mostly by radiation or chemical mutagenesis and are of great interest in the mushroom industry as they alleviate problems caused by spore formation^10, 12, 46, 47^. We anticipate that *Srr1* or its target genes could be targets of breeding novel sporeless strains of commercially cultivated species, such as *Pleurotus* spp or *Lentinula edodes*.

To reconstruct genes downstream of SRR1, we performed RNA-Seq on the deletion mutant. Genes that failed to be activated in the mutant compared to the wild-type included groups related to the fungal cell wall, aromatic amino acid biosynthesis, TOR signalling and secreted proteins, among others. The promoters of the 559 downregulated genes provided a sharp signal for a binding motif of SRR1 which is supported by an independent, protein sequence-based predictor, allowing the inference of direct and indirect target genes, even in the absence of ChIP-Seq, which is not currently available for *C. cinerea*. Based on the available information, we illustrate a putative regulatory network in which SRR1 serves as the central regulator, linking upstream factors and downstream targets involved in spore morphogenesis. It should be emphasised that although there was a sharp signal for the SRR1 motif in the RNA-Seq data, this network is derived from indirect evidence on regulatory interactions, thus requiring careful interpretation. Nevertheless, the hypothetical regulatory network of SRR1 reveals potential interactions, pathways, other transcription factors and target genes activated during spore morphogenesis.

Ballistospory is one of the most defining traits of the Basidiomycota. We reasoned that, as spore formation is a terminal process in mushroom development, genes defining ballistospory should be downstream of SRR1 and that they should be missing in secondarily non-ballistosporic Agaricomycetes (e.g. puffballs, false-truffles). Using phylostratigraphy of SRR1 downregulated DEGs and comparative genomics, we identified twelve genes that show links to ballistospory. Analysis of one of these, the chitinase *ChiE2* showed that its deletion causes a spore-poor phenotype and an abnormal spore shape with a partial loss of asymmetry. As the latter is a key trait in ballistospory and ChiE2 orthologs are lost in secondarily non-ballistosporic species, we speculate it may be involved in this process. However, it is also plausible that secondarily non-ballistosporic taxa lost ChiE2 for reasons other than forcible spore discharge. In addition to *ChiE2*, further genes are certainly required for ballistospory. Our phylostratigraphic analyses revealed some that could speculatively be linked to certain steps in this process (e.g. transporters for the growth of Buller’s drop), though evidence for these is preliminary and they require further research. Thus, although our search did not provide a comprehensive answer for the genetics of ballistospory, it did provide the first insights into this unique process and we anticipate that future analyses of SRR1 target genes should reveal further genes involved.

Taken together, this study has provided novel insights into the genetics of postmeiotic spore morphogenesis and, to some extent, the unique spore discharge mechanisms of the Agaricomycetes. While the genetics of these processes is certainly more complex than what we could decipher in this study, we anticipate that our results will contribute to understanding the general principles of sexual spore dispersal in a large and diverse group of fungi and will inspire further research on spore morphogenesis, ballistospory and mushroom biotechnology.

## Acknowledgements

We acknowledge support by the ‘Momentum’ program of the Hungarian Academy of Sciences (contract no. LP2019-13/2019 to L.G.N.) the European Research Council (grant no. 101086900 to L.G.N.), the Eötvös Loránd Research Network (SA-109/2021, to L.G.N.) as well as the National Key Research and Development Program of China (2022YFD1200600, to W.G.). Z.H. acknowledges the support of China Scholarship Council (grant no. 202008110168) and Stipendium Hungaricum Scholarship (grant no. SHE-21070-006/2020).

## Author contributions

Z.H. and L.G.N. conceived the study. Z.H., Y.Y., Y.Z, A.C., C.F., H.W., Z.K., E.A., N.M., M.V., X.B.L., N.Z., and I.K. performed the wet-lab experiments. Z.H., Z.M., B.B., and B.H. processed the bioinformatic data. Z.H., Z.M., Y.Y., L.G.N., and W.G. analysed the data and prepared figures. Z.H., L.G.N., Z.M., and W.G., wrote the original manuscript. All authors have read and commented on the manuscript.

## Materials and Methods

### Gene expression datasets

The raw RNA-Seq data of different developmental stages in *Coprinopsis cinerea* was obtained from Krizsán et al.^25^, and reanalysed using the reference genome and the reanalysis workflow by Botond et al.^34^. The raw expression data of *Pleurotus ostreatus, Laccaria bicolor* and *Cyclocybe aegerita* were obtained from the published developmental transcriptomes^26, 27, 28^, and the orthology relationships of SRR1 were generated from the results of Nagy et al.^43^. Gene expression levels were shown in the line charts and expression fold-changes were shown with a barplot, plotted with the R package ggplot2^48^.

### Strains and culture media

The homokaryotic *C. cinerea A43mut B43mut pab1-1* #326 strain and dikaryotic *P. cornucopiae* CCMSSC00406 strain (from China Centre for Mushroom Spawn Standards and Control) were used in this study. For oidiation of *C. cinerea,* small agar plugs were inoculated on yeast extract-malt extract-glucose (YMG) agar medium and cultured at 37 °C for 6-7 days under constant light to generate the oidia for the transformation. For fruiting *C. cinerea* on the Petri dish, the strains were grown on YMG (with 2% glucose) medium at 28 °C for 5-6 days until the mycelium reached ∼1 cm of the edge of the Petri dish, then were transferred to white light for 2 h and back to dark for another 1 day, followed by a 12/12 h light/dark cycle for fruiting, at 28 °C with around 80% relative humidity. For phenotyping *C. cinerea* on straw, the substrate was prepared as follows: 75% straw and 25% wheat bran, with the water content around 80%. For *P. cornucopiae*, the Potato Dextrose Agar (PDA) medium was used for the basic culturing, and the substrate as follows was used for fruiting: 94% cottonseed hulls, 5% wheat bran, and 1% gypsum, with an approximate water content of 65%. Then polycarbonate plastic bottles (diameter=70 mm, height=90 mm, volume=280 ml) were filled with around 180 g of substrate and autoclaved for 2 hours at 0.11 MPa and 121 °C.

### Plasmid construction, transformation, RNA interference and the mutant screening

For *C. cinerea*, the CRISPR/Cas9 system was used for gene deletion. The primers and crisprRNAs (crRNAs) are listed in the Supplementary Table 6. The backbone was linearized by the primer pair PUC19_F/PUC19_R for *Srr1*, and by primer pair PUC19_F_1/PUC19_R_1 for *ChiE2* using pUC19 plasmid as a template. The native para-aminobenzoic acid (PABA) synthase gene (*pab1*) was amplified from the pMA412 vector using the primer pair Srr1_PABA_1F/Srr1_PABA_1R (for *Srr1*) and primer pair PABA_F/PABA_R (for *ChiE2*) as the positive selection marker^49, 50^. The upstream and downstream homology arms of *Srr1* were amplified by the primer pairs Srr1_Up_1F/Srr1_Up_1R, and Srr1_Down_1F/Srr1_Down_1R, respectively, using genomic DNA as a template. The fragments were assembled by Gibson Assembly Cloning Kit (NEB, UK) according to the manufacturer’s instructions. The assembled plasmid map is shown in Supplementary Fig. 2A for *Srr1* and Supplementary Fig. 11A for *ChiE2*. Two sgRNAs (srr1_sgRNA_1 and srr1_sgRNA_2) were designed for *Srr1* and one (ChiE2_sgRNA) for *ChiE2* by sgRNAcas9^51^. The *in vitro*-assembled RNP complexes were prepared using commercially synthesized trans-activating CRISPR RNA (tracrRNA) and CRISPR RNA (crRNA) according to the manufacturer’s instructions (IDT, USA). For each *in vitro*-assembled RNP mixture, 1.2 μL of crRNA (100 μM), 1.2 μL of tracrRNA (100 μM) and 9.6 μL of duplex buffer were mixed. The duplexes were incubated at 95°C for 5 minutes, followed by cooling for 2 minutes. The RNA duplexes, along with Cas9 (10 μg/μL), 1.5 μL Cas9 working buffer, and 1.0 μL duplex buffer were then mixed and incubated at 37 °C for 15 minutes. Transformation of *C. cinerea* was conducted according to the protocol described by Dörnte et al. and Pareek et al.^29^ ^52^. The DNA samples for screening transformants were prepared with the microwave-based method^53^. For *Srr1*, the primer pair Srr1_Up_1F/Srr1_Down_1R was used for the first-round screening, samples with products in different sizes from WT (3906 bp) were selected for the second round of screening. The primer pair Srr1_GF/Srr1_GR was used for the second round of screening, and transformants with no WT band (1365 bp) were selected as clean mutants (Supplementary Fig. 2B-D). RT-PCR was also performed to verify the lack of transcription of Srr1 using the primer pair Srr1_RT_1F/Srr1_RT_1R. For *ChiE2,* the primer pair ChiE2_CF/ChiE2_CR was used for PCR screening, because the plasmid can be multiple or partially integrated in the cleavage site, samples with missing (too long to amplify) or different sized (partially integrated) product from WT (845 bp) were selected as the mutants (Supplementary Fig. 11B-C).

For *P. cornucopiae*, the RNA interference (RNAi) method was conducted for the gene knockdown. The primers are listed in the Supplementary Table 6. An RNAi plasmid was constructed using the modified pCAMBIA1300 vector containing the hygromycin B phosphotransferase gene (*hyg*)^54^. The RNAi-F and RNAi-R sequences were PCR amplified using genomic DNA from strain CCMSSC00406 as a template and then ligated into the interference vector (Supplementary Fig. 4). Subsequently, the RNAi plasmid was transferred into the wild-type strain using *Agrobacterium tumefaciens* GV3101, following previous research protocol^55^. Two rounds of selection of transformants were performed on complete yeast medium (CYM) supplemented with 90 mg/ml hygromycin B. Transformants with the highest interference efficiency were screened through quantitative PCR (qPCR) analysis. RNA extraction was performed using the E.Z.N.A. Plant RNA Kit (Omega Bio-Tek, Norcross, GA, USA), and cDNA was synthesised using the HiScript Q RT SuperMix reverse transcription kit (Vazyme, Nanjing, China). qPCR reactions were carried out using AceQ qPCR SYBR Green Master Mix (Vazyme, Nanjing, China) and an ABI 7500 real-time PCR amplifier (Applied Biosystems, Foster City, CA, USA). The amplification program included an initial denaturation at 95°C for 30 seconds, followed by 40 cycles of 95°C for 10 seconds, 60°C for 30 seconds, and extension at 72°C for 30 seconds. The 2^−ΔΔCT^ method was used to calculate the relative gene expression levels, with the *β-actin* gene used as a reference.

### Phenotyping of *C. cinerea*

For growth rate measurement, small agar plugs (d=5 mm) of wild-type and Δ*Srr1* strains (3 replicates/group) were inoculated on the YMG (2% glucose) media and cultured at 28 °C in the dark. Colony diameters were measured every 24 h until 96 h. For testing the oidiation, wild-type and Δ*Srr1* strains (3 replicates/group) were cultured on YMG (2% glucose) media at 37 °C under constant light for 4 days. The colony diameters were measured and the oidia were collected, counted and quantified by oidium number/mm^2^. For analysis of fruiting body development, the strains were inoculated on the straw substrate and grown at 37 °C in the dark. After the mycelium was fully grown, they were transferred into the incubator with a 12/12 h light/dark cycle at 28 °C with ∼80% relative humidity for fruiting. Photos of fruiting bodies were taken in different developmental stages and the time-lapse videos were generated for monitoring the fruiting process. For basidiospore counting, the whole mature caps (after the fruiting body autolysis) were collected (7 for WT and 8 for Δ*ChiE2*), weighed and then were put into the 1.5ml Eppendorf tube with 500 μL water. The caps were smashed by the small pestle to detach the basidiospores. The basidiospores were counted and quantified by the number of spores/gram cap, the length and width of the spores (20 for each strain) were measured and the aspect ratio (length/width) was calculated. For data illustration, the line charts and the boxplots were plotted in R with the ggplot2 package^48^, and the Student’s t-test was used to assign the significance with the ggsignif package^56^.

For checking the meiosis in Δ*Srr1* and WT by microscopy, we took gill samples of WT and Δ*Srr1* fruiting bodies when spore initials emerge at the sterigma tips (the Δ*Srr1* arrest stage). Hoechst 33342 (0.1 μg/ml, Thermo Fisher Scientific, UK) was used for staining nuclei for 10 minutes, followed by 2 times washing using PBS buffer. The samples were checked under a fluorescence microscope both under bright and the blue channel. For comparing the morphology of basidia and basidiospores in WT and Δ*ChiE2*, we took the samples from the gill of the WT and Δ*ChiE2* fruiting bodies around the sterigmata formation and spore maturation stages (according to the definition of Kues^40^), and checked them under the optical microscope.

### Phenotyping of *P. cornucopiae*

The *P. cornucopiae* strain was pre-cultured on the PDA media at 28 °C for 7 days in the dark. Well-grown agar plugs (3 pieces) were taken from PDA plates and inoculated into the substrate in the plastic bottles, followed by cultivation in the dark at 25 °C. After ∼15-20 days. The bottles were opened when the mycelium fully filled the substrate, then the surface-aged mycelium was removed, and the plastic bottles were transferred to environmental-controlled fruiting facilities at 18 °C. Relative air humidity was maintained at 85%-95%, light intensity was set at 300-500 lux with a daily illumination period of 12 hours, ventilation was performed 3-4 times, and CO_2_ concentration was kept below 1000 ppm. Each experimental group included 20 replicates.

Spore quantification was conducted when the fruiting bodies reached maturity. Mature mushroom caps were carefully placed on glass slides, were let to drop spores, then covered with a coverslip, and observed under a microscope, with photographs taken. Mature mushroom caps were also placed on black paper, covered tightly, and after 12 hours, spore prints were obtained. A 0.5 x 0.5 cm portion was cut from the edge of the obtained spore print (where spores were most concentrated), rinsed thoroughly with 500 μL ddH_2_O, resulting in a spore suspension. The suspension was then diluted tenfold, and spore counting was performed using a hemocytometer.

The formula for spore count per ml is given by:

Spore count (per ml) =Total number of spores in 80 small squares/(80×400×10000×10) This calculation involves counting spores in 80 small squares, where each small square corresponds to a specific volume and dilution factor.

### RNA-sequencing and data analysis

For collecting samples for the RNA-Seq, the gills were isolated by peeling cap and veil tissues off of gills using a scalpel and forceps from the Δ*Srr1* and WT young fruiting bodies (3 replicates for each group) when spore initials emerge at the sterigma tips (Fig. 4A), and put in the liquid nitrogen immediately. RNA extraction was performed using the Quick-RNA Miniprep Kit (Zymo Research, USA), non-strand-specific cDNA library and pair-end (150 bp) sequencing were conducted by Novogene Co., Ltd (Cambridge, UK) with a range of 41-54 million reads per sample. The raw reads were checked by FastQC (version 0.11.8),^57^ and trimmed by bbduk.sh (version 38.92, qtrim=r trimq=10 minlength=20)^58^. Trimmed reads were mapped to the reference genome using STAR (version 2.6.1, --alignIntronMin 20 -- alignIntronMax 3000)^59^. Featurecount (version 2.0.1) was used for quantification based on CDS features in the GFF file^60^.The raw count after Variance Stabilizing Transformation (VST) by the DEseq2 package was used for sample clustering by Pearson correlations^61^, and the global similarity was visualised by the R package ComplexHeatmap^62^.The genes that have more than five raw counts in at least two replicates in one group were kept. For identifying the differentially expressed genes (DEGs), the R package edgeR was used^63^. Genes with a fold-change ≥ 2 and adjusted p-value (Empirical Bayes Statistics) ≤ 0.05 were considered as upregulated DEGs, while genes with an adjusted p-value (Empirical Bayes Statistics) ≤ 0.05 and fold-change ≤ −2 and were considered as downregulated DEGs. The volcano plot was generated by the R package EnhancedVolcano^64^. The gene expression levels of downregulated DEGs of different stages were obtained from Krizsán et al.^25^, and the expression heatmap was generated by R package ComplexHeatmap with row scale^62^.

The secreted proteins were predicted by SignalP (version 6.0) with default parameters^65^. The InterPro annotation was performed by InterProScan (version 5.65-97.0) with ‘--goterms’ parameter to map the InterPro annotation to the GO terms^66^, and the GO enrichment was performed by R package clusterProfiler^67^, the p-value cut-off was set to 0.05, and the minimal size of genes annotated for testing was set to minGSSize=2. For the KEGG enrichment, we first found the reciprocal best hits (RBH) of the downregulated genes between *C. cinerea* Amut1Bmut1 and *C. cinerea* Okayama 7 using the “easy-rbh” module of MMseqs2 (version 13.45111)^68, 69^. Then the KEGG enrichment of the downregulated genes was conducted by R package clusterProfiler using *C. cinerea* Okayama 7 (ID: cci) KEGG genes database^67^, the p-value cut-off was set to 0.05, and the minimal size of genes annotated for testing was set to minGSSize=2.

### Identification of downstream targets and construction of regulatory network

To infer the binding motif of SRR1, six promoter datasets were established based on different upstream (or intron) regions of down-regulated DEGs (Fig. 5A). We extracted the 300 bp and 500 bp upstream sequences of the start codon for the fold-change two downregulated DEGs (datasets: FC2_upstream_300bp and FC2_upstream_500bp, respectively). We also isolated the complete upstream intergenic region including the first intron from fold-change two downregulated DEGs (dataset FC2_intergenic_first_intron). Similarly, three datasets of fold-change four downregulated DEGs were also created, named FC4_upstream_300bp, FC4_upstream_500bp, FC4_intergenic_first_intron. Analyses of motif enrichment were carried out in STREME with the following parameters: minimum motif width 6 bp, maximum motif width 10 bp, p-value cut-off 0.05, other parameters were set to default^70^.

Another independent motif prediction was performed by the online tool (https://zf.princeton.edu/) using the protein sequence of SRR1^42^. Genes with the inferred motif in FC2_intergenic_first_intron dataset were considered as the direct targets of SRR1 while others were considered as the non-direct targets. The motif distribution was illustrated for the direct targets in the FC2_intergenic_first_intron dataset with TBtools (version 2.080)^71^. The regulatory network was visualized by Cytoscape (version 3.10.2) with the direct/non-direct regulatory relationships and functional group annotation^72^.

### Phylogenetic and phylostratigraphic analyses

For the species tree, we selected 189 species representing the Basidiomycota with Ascomycota as outgroups. The protein files of these species were downloaded from JGI (04/2023). We used the same set of marker genes as in Merényi et al.^26^; these were identified in a reciprocal best hit (RBH) search between *C. cinerea* (CopciAB) and all proteins of the 189 species. This method was used to construct 177 clusters of single copy genes, which were aligned using PRANK (version 170427)^73^, and trimmed using the TrimAL (version 1.2rev59) “-strict” approach^74^. For the supermatrix alignments with a length of at least 100 amino acids and presence of at least 75 species were required. Species tree reconstruction was performed using maximum likelihood inference (ML) in IQ-TREE (version 1.6.12)^75^. For each partition, the LG+G substitution model was applied and 1,000 ultrafast bootstrap replicates were used to assess branch support.

Phylogenetic analyses of SRR1 orthologs were carried out with first identifying homologs identified through protein BLAST with an e-value cutoff of 10^−10^ in the genomes of 189 fungi. Retained homologs were aligned by mafft (version 7)^76^, followed by manual inspection during which overly short (<50 aa) or divergent sequences were removed. Representatives of the most closely related C2H2-type zinc finger protein subfamily were used as an outgroup. Phylogenetic analysis was carried out in RAxML (version 8.2.12) under the CAT + WAG model with 500 rapid bootstrap replicates^77^.

For the phylostratigraphic analysis, we used the reciprocal best hit approach mentioned above. Based on the species content of these clusters, their origin was mapped to the corresponding ancestors using a custom R script.

To identify *Coprinopsis* genes that might be relevant for ballistospory, we combined three data sets. 1) RBH search between *C. cinerea* and 189 genomes mentioned above 2) Developmental transcriptome from fruiting body formation of *Coprinopsis*^25^, and 3) Developmental transcriptome of eight Agaricomycotina species analysed in Nagy et al.^43^. We used Dataset 1 to characterise gene conservation among the Basidiomycota and to identify losses among the secondarily non-ballistosporic species. For screening candidate ballistospory-related genes we required 50% presence (RBH hits) among ballistosporic species, losses in at least 5 of the 8 independent secondarily non-ballistosporic clades, but at most 30 losses across the whole species tree. Dataset 2 was used to confirm the lamella-specificity of *Coprinopsis* genes. We required the highest expression in lamellae. In Dataset 3, tissue-specific expression of 1:1 orthologs was inferred by first calculating a tissue-specificity score following et al.^78^. For tissue-specific expression we required tissue-specificity score ≥1 and at least four 1:1 orthologues to show lamellae or cap (because lamellae samples were not available in each species) specific expression.

