## Supplementary Figures for "A new regulator of sporulation sheds light on spore morphogenesis and ballistospory in mushroom-forming fungi"

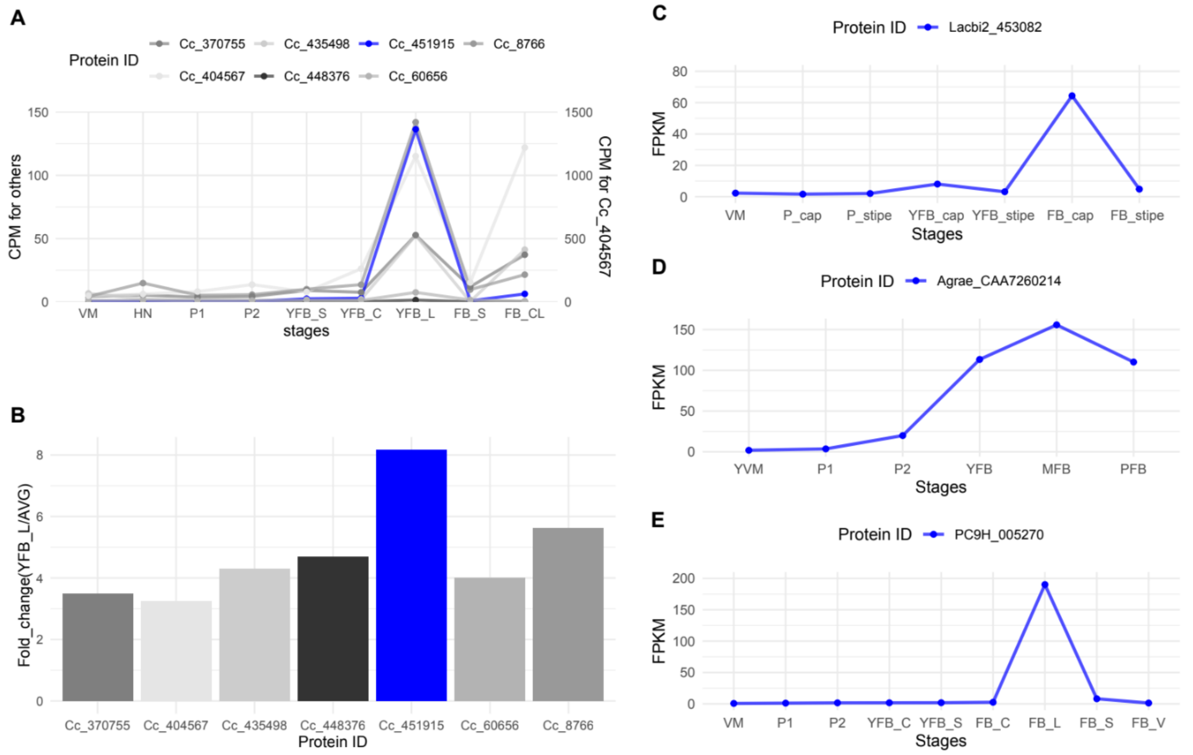


***Supplementary Figure 1. Expression levels of gill-upregulated transcription factors in different mushroom-forming fungi.*** *A. The expression level of seven gill-upregulated transcription factors in Coprinopsis cinerea across developmental stages[1]. VM: vegetative mycelium, H: hyphal knot, P1: primordium stage 1, P2: primordium stage 2, YFB_S: young fruiting body stipe, YFB_C: young fruiting body cap,YFB_L: young fruiting body lamella, FB_S: mature fruiting body stipe, FB_CL: mature fruiting body cap and lamella. B. The fold-change of the expression level of Cc_451915 (Srr1) in young fruiting body cap compared to the average expression level of all stages in C. cinerea. C. The expression level of Srr1 ortholog in Laccaria bicolor (Lacbi2_453082) across developmental stages[2]. VM: vegetative mycelium, P_cap: primordium cap, P_stipe: primodium stipe, YFB_cap: young fruiting body whole cap, YFB_stipe: young fruiting body stipe. FB_.cap: fruiting body whole cap, FB_stipe: fruiting body stipe. D. The expression level of Srr1 ortholog in Cyclocybe aegerita (Agrae_CAA7260214) during the different developmental stages[3]. YVM: young vegetative mycelium, P1: primodia stage 1, P2: primordium stage2 YFB: young fruiting body, MFB: mature fruiting body. PFB: post sporulating fruiting body. E. The expression level of Srr1 ortholog in Pleurotus ostreatus across developmental stages[4]. VM: vegetative mycelium, P1: primordium stage 1, P2: primordium stage 2, YFB_C: whole young fruiting body cap, YFB_S: young fruiting body stipe, FB_C: fruiting body cap, FB_L: fruiting body lamella, FB_S: fruiting body stipe, FB_V: fruiting body cuticle.*

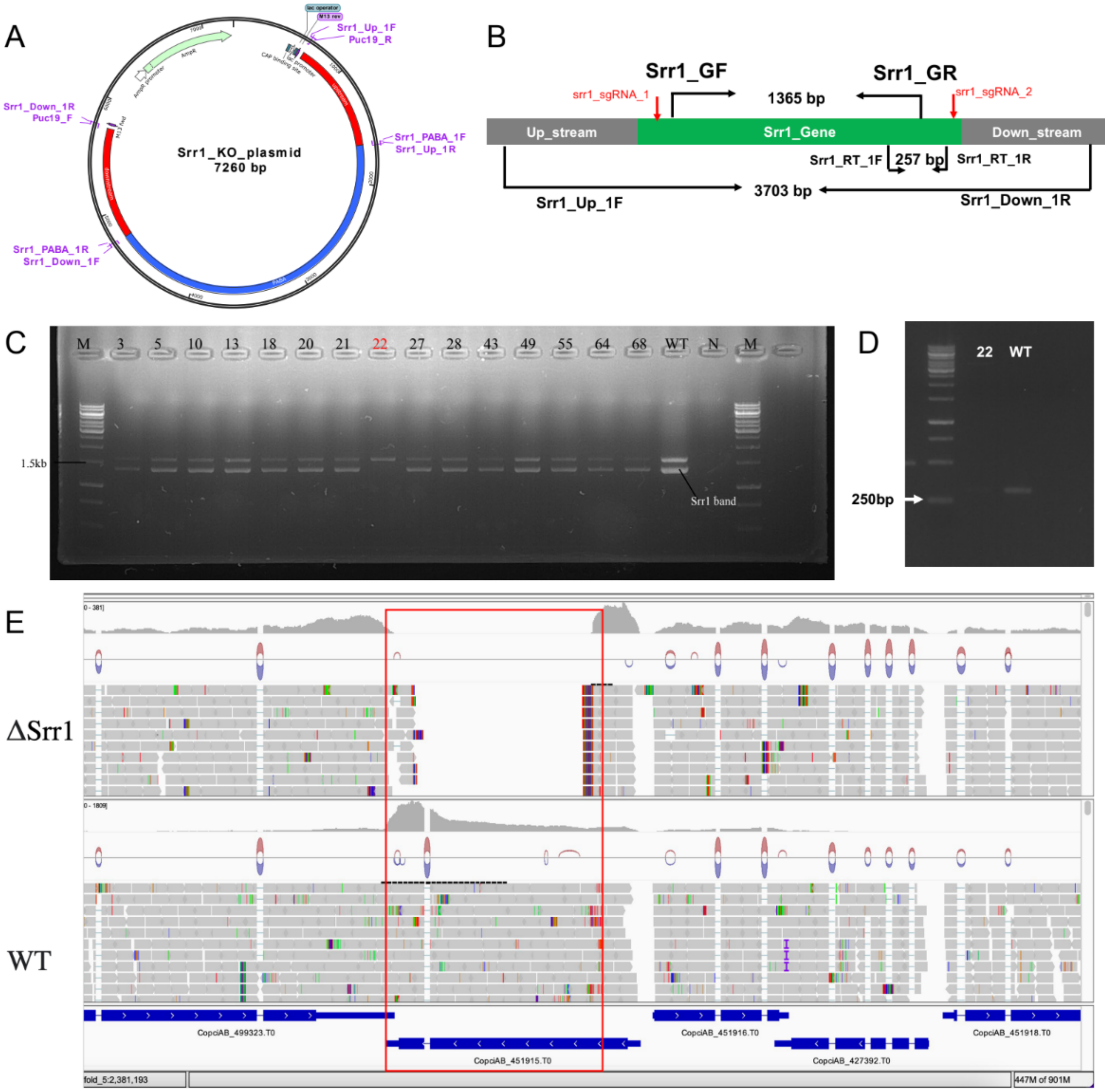


***Supplementary Figure 2. The plasmid map and screening result of the Srr1 deletion.*** *A. The map of the plasmid used for knock-out. B. Schematic diagram showing the position of screening primers and single-guide RNA. C. Agarose gel electrophoresis result of PCR-based screening of positive transformants (highlighted in red) using Srr1_GF/ Srr1_GR primer pair. D. The gel electrophoresis of RT-PCR results using the srr1_RT_1F/srr1_RT_1R primer pair. E. IGV visualization of the mapped reads of wild-type (WT) and ΔSrr1 in the RNA-Seq data.*

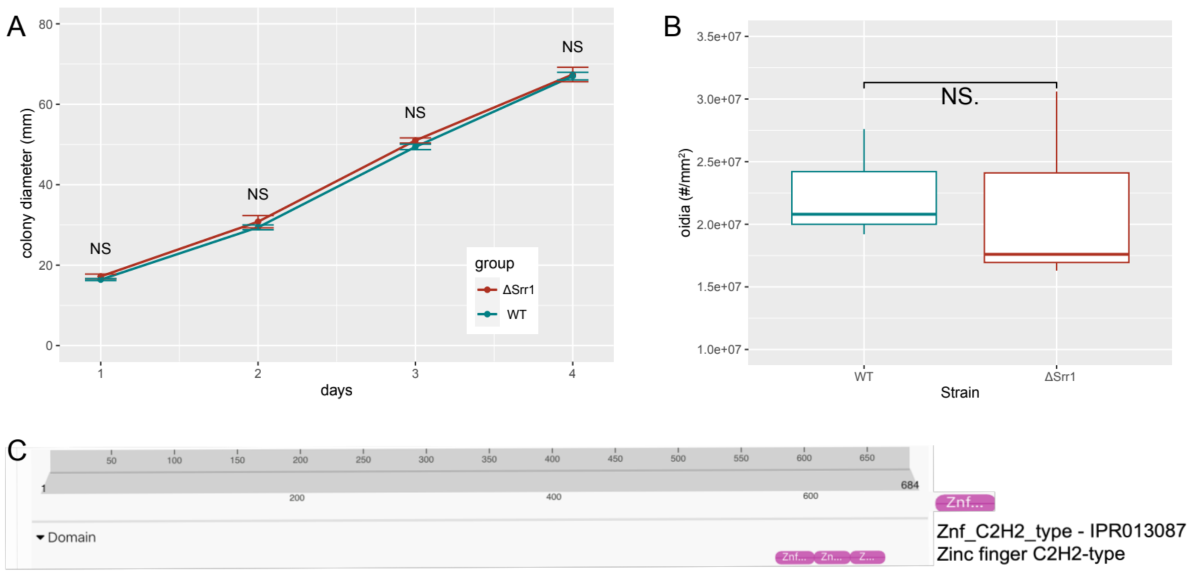


***Supplementary Figure 3. The comparison of growth rate, oidium quantification in wild-type (WT) and ΔSrr1, and the InterPro annotation of SRR1.*** *A. The line chart showing the vegetative mycelium growth rate of WT and ΔSrr1. The error bars represent the standard deviation of three replicates, the Student's t test is used to compare the means between two groups, NS: no significance. B. The boxplot showing the count of oidia/mm$^2$ in WT and ΔSrr1. Each group has three replicates, the Student's t test is used to compare the means between two groups, NS: no significance. C. The InterPro annotation results of the SRR1 protein.*

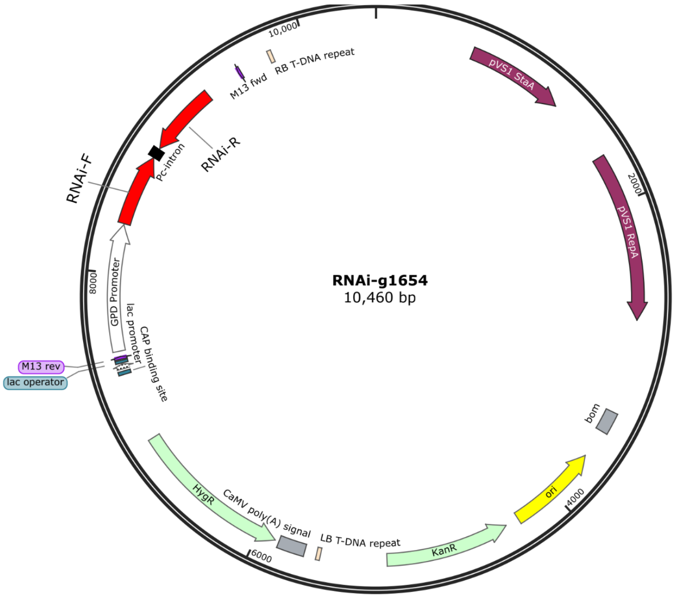


*Supplementary Figure 4. The map of the RNA interference plasmid for P. cornucopiae Srr1 (g1654).*

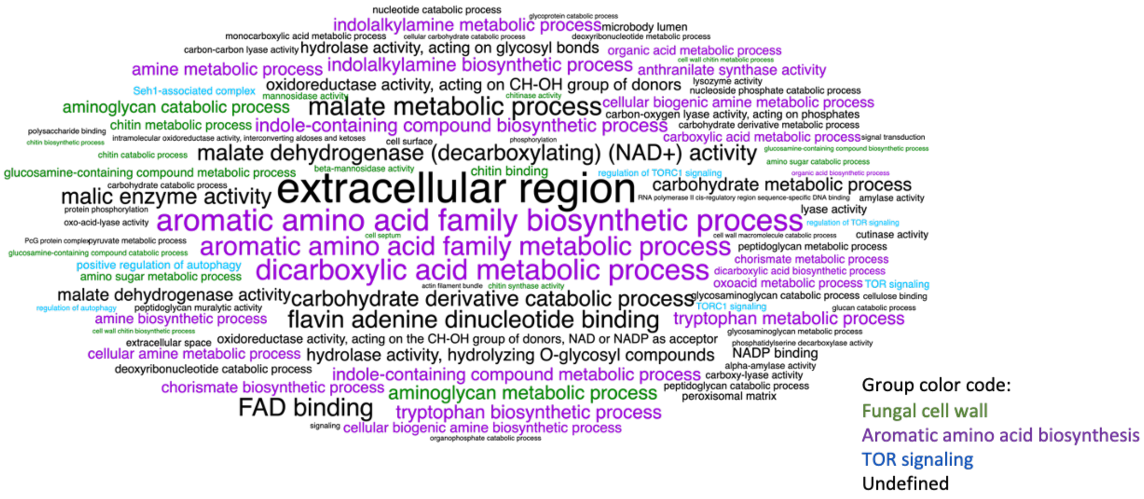


***Supplementary Figure 5. A word cloud of the enriched GO terms of downregulated DEGs (Supplementary Table 2).*** *The word frequency (size of the word) is based on -log*$_{10}$*(p-value) and the colour is based on the different functional groups.*

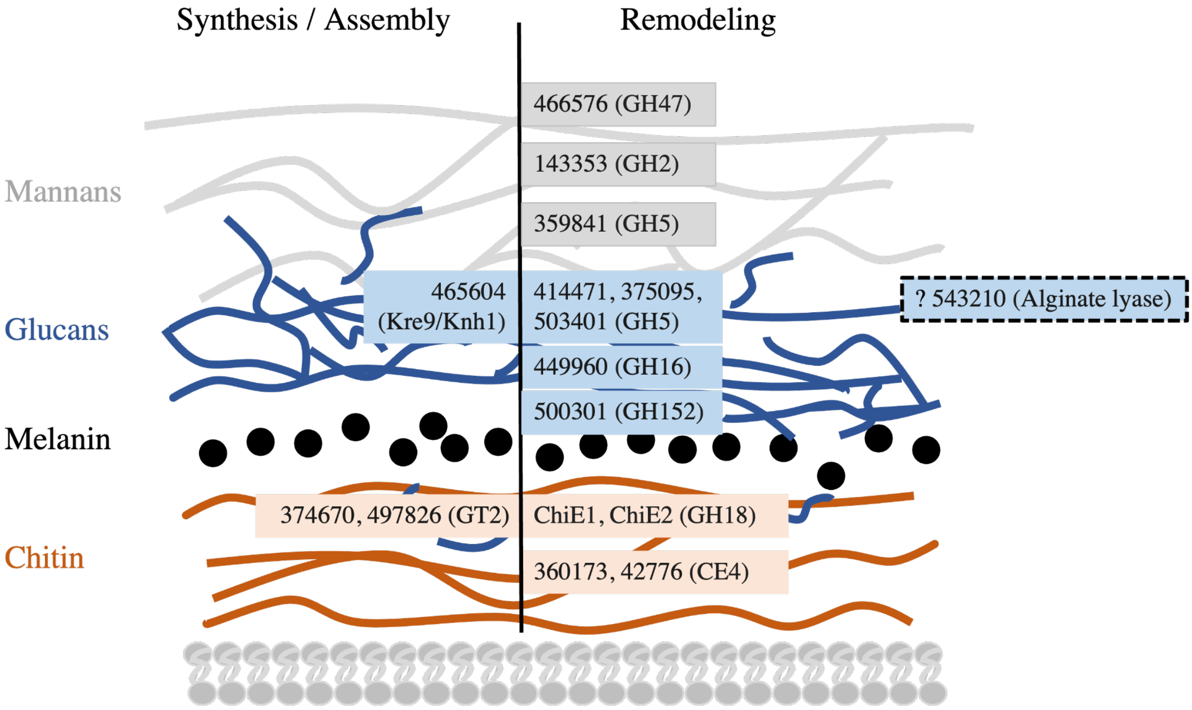


***Supplementary Figure 6. Schematic diagram of the spore wall in C. cinerea with the putative related downregulated genes, adapted from Ehren et al.* [5]*.*** *The melanin layer only represents its presence, but the position of the layer is unknown. Boxes with dotted borders represent the prediction with less evidence.*

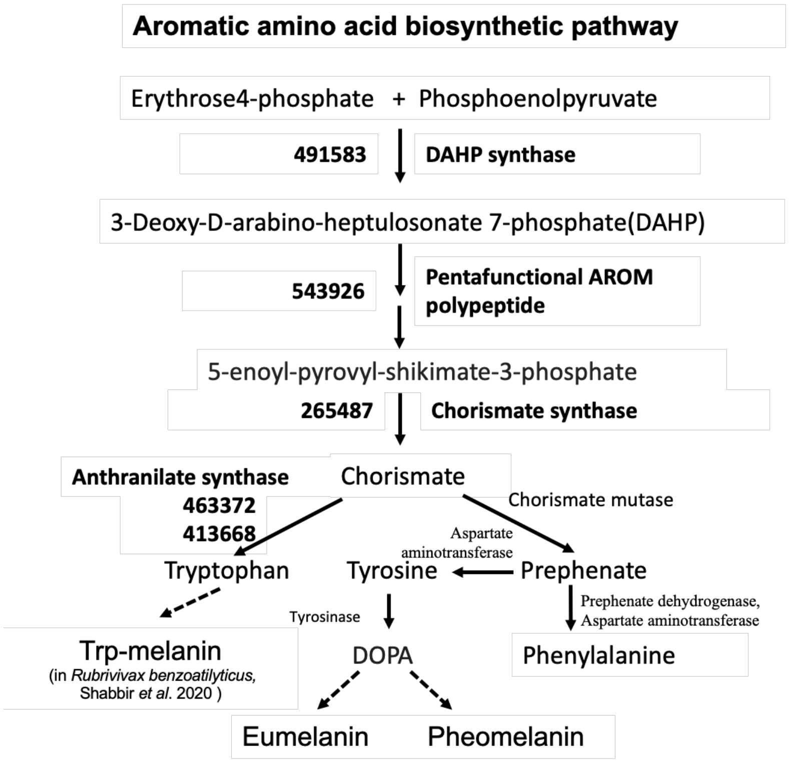


***Supplementary Figure 7. Diagram of the aromatic amino acid biosynthesis based on the KEGG pathway (ID: cci00400).*** *Gene IDs and names with bold text represent downregulated DEGs in ΔSrr1. The dotted arrows represent unclear parts of the pathways in basidiomycetes.*

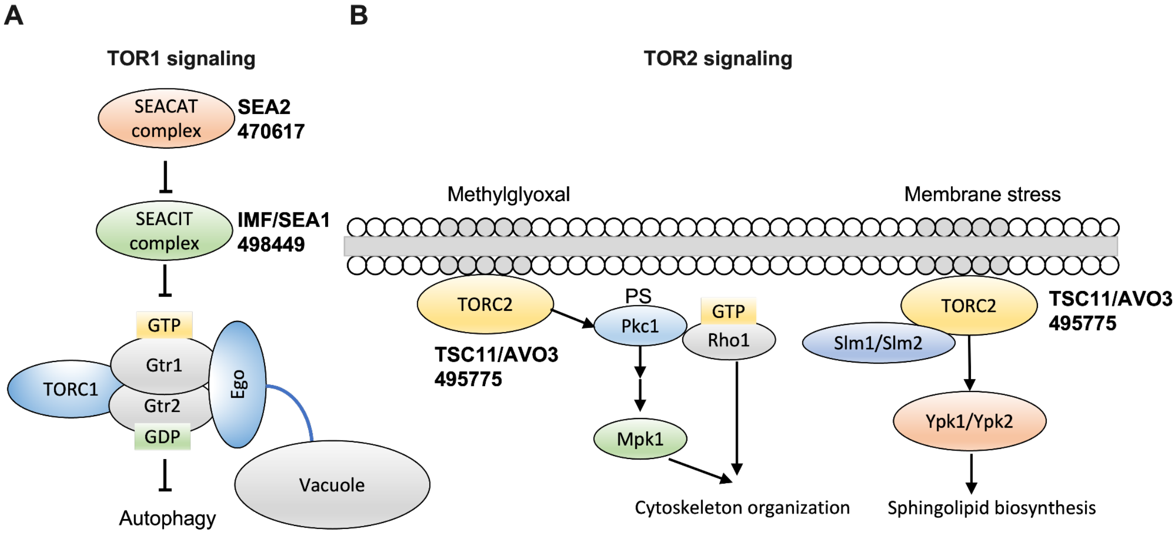


***Supplementary Figure 8. The schematic diagram of TOR signalling pathway, adapted from the pathways in yeast[6].*** *A. TOR1 signalling pathway. B. TOR2 signalling pathway. Genes with bold text represent downregulated DEGs in ΔSrr1.*

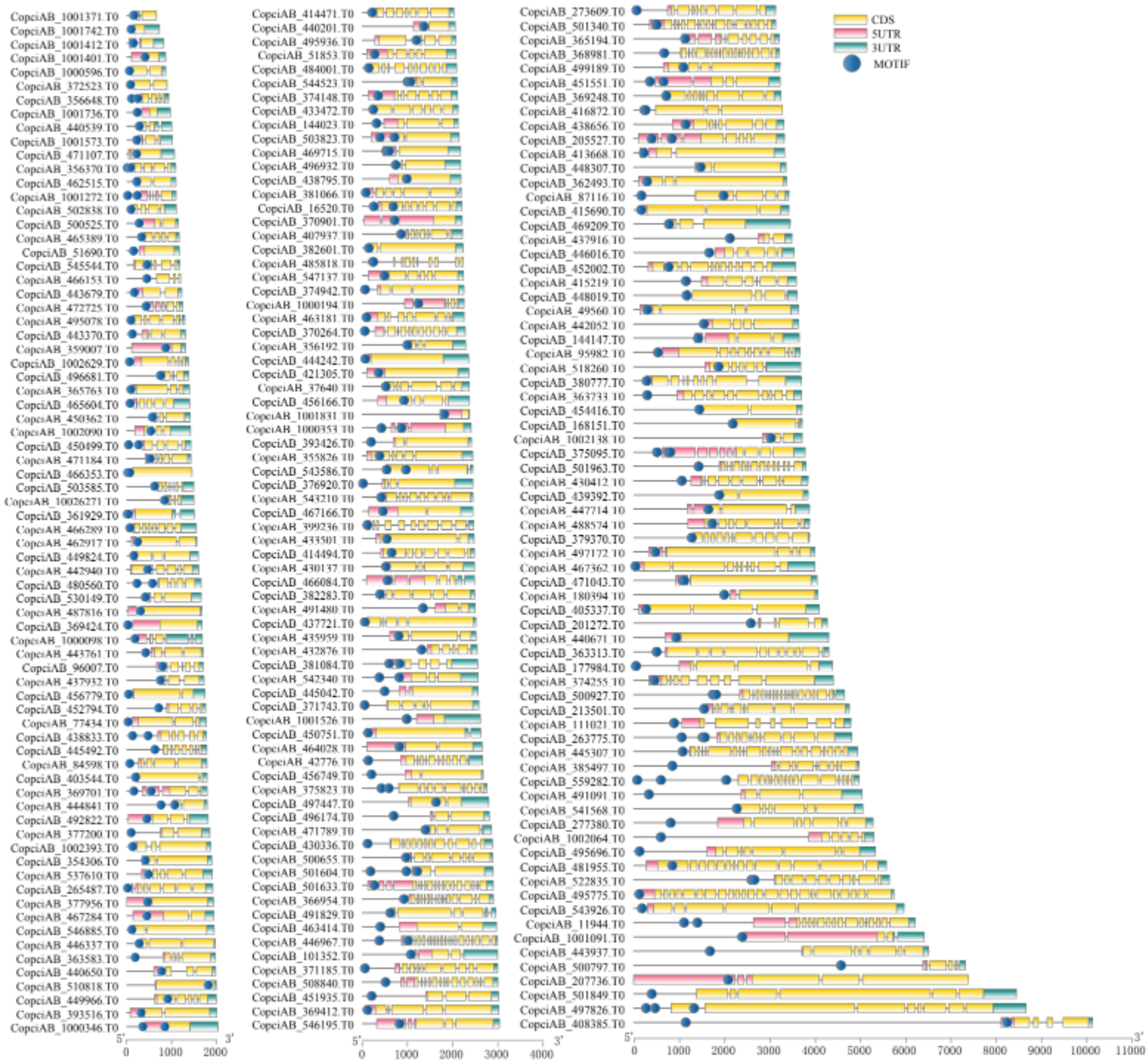

***Supplementary Figure 9. The distribution of the inferred motif in the complete upstream intergenic region (including the first intron) from downregulated DEGs (Down_FC2_Intergenic_first_intron dataset).***

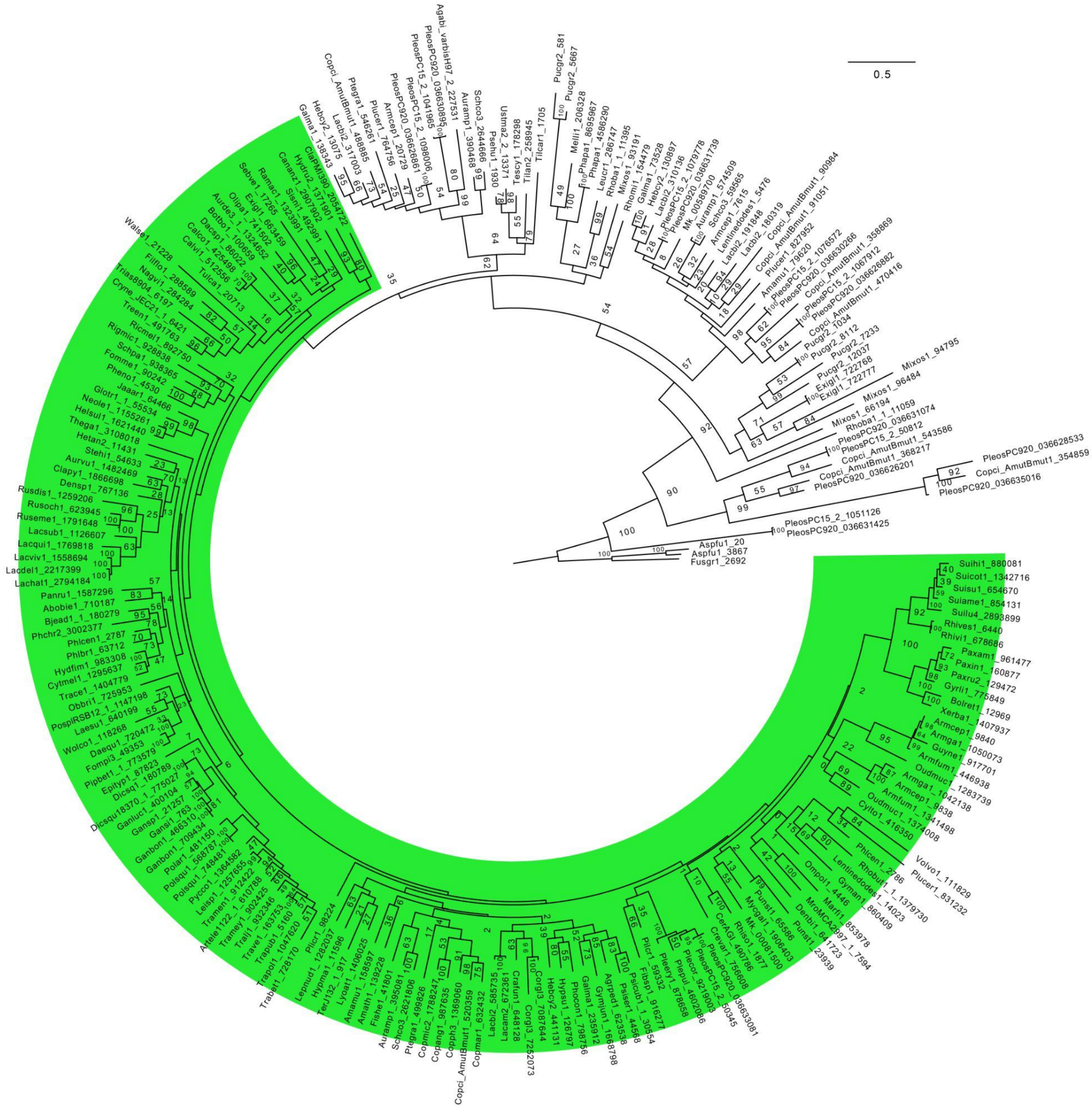


***Supplementary Figure 10.*** *Phylogenetic and orthology relationships of ChiE2 across the Agaricomycetes. The green highlighted clade contains orthologs of C. cinerea ChiE2. The tree is based on Maximum Likelihood, inferred under the PROTGAMMAWAG model of evolution. Bootstrap values are shown next to branches.*

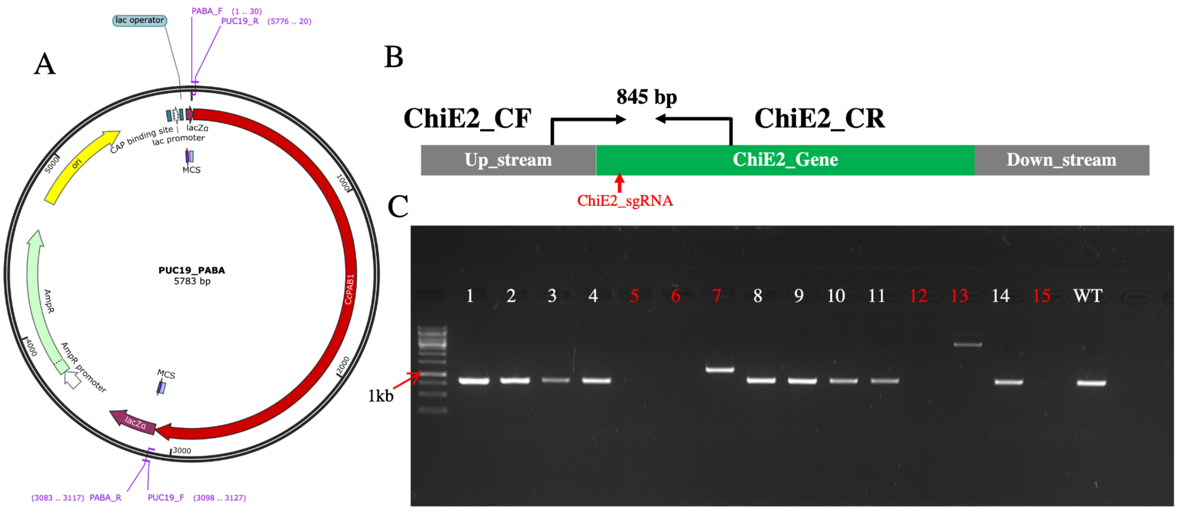


***Supplementary Figure 11. Knockout of the ChiE2 gene**. A. The map of the plasmid used for knock-out. B. Schematic diagram showing the screening primers and single-guide RNA. C. Agarose gel electrophoresis result of PCR-based screening of positive transformants (highlighted in red) using ChiE2_CF/ChiE2_CR primer pair.*

6. Yoshiharu I, Wataru N. TOR Signaling in Budding Yeast. In: *The Yeast Role in Medical Applications* (ed Waleed Mohamed Hussain A). IntechOpen (2017).
